# Nutritional regulation of cellular quiescence depth and cell cycle re-entry in Vasa2+/Piwi1+ cells in a sea anemone

**DOI:** 10.1101/2025.02.27.640509

**Authors:** Eudald Pascual-Carreras, Kathrin Garschall, Patrick R.H. Steinmetz

## Abstract

Animals with lifelong growth adjust their growth rates to nutrient availability, yet the underlying cellular and molecular mechanisms remain poorly understood. Here, we studied how food supply and TOR signalling regulate the cell cycle in a multipotent, Vasa2-/Piwi1-expressing cell population in the sea anemone *Nematostella vectensis*. We discovered that starvation induces a reversible G_1_/G_0_ cell cycle arrest in Vasa2+/Piwi1+ cells and that cell cycle re-entry upon refeeding is dependent on TOR signalling. In addition, the length of the refeeding stimulus after starvation determines the proportion of cells that re-enter S-phase. Remarkably, prolonged starvation delayed both refeeding-induced TOR signalling activation and S-phase re-entry. This strongly suggests that *Nematostella* Vasa2+/Piwi1+ cells undergo starvation-controlled quiescence deepening, previously described only in unicellular eukaryotes and mammalian cell culture. The nutritional control of quiescence and cell proliferation may thus be a fundamental, evolutionarily conserved strategy underlying the environmental regulation of indeterminate growth in animals.

## Introduction

Animals with indeterminate growth retain the ability to regulate their body growth throughout life in response to changing environmental conditions. They are found across diverse lineages such as fish, crustaceans, annelids and most non-bilaterian taxa including sea anemones, corals and sponges (Hariharan et al., 2015; Leys and Lauzon, 1998; Sebens, 1987). Lifelong growth and high body plasticity are therefore likely ancestral to all animals (Hariharan et al., 2015; Vogt, 2012). Some animals with indeterminate growth dramatically adjust their body size to environmental changes, shrinking during starvation and regrowing upon refeeding. Examples of whole-body shrinkage include marine iguanas (Wikelski and Thom, 2000), planarians (Baguñà et al., 1990) and many cnidarians such as sea anemones (Garschall et al., 2024), hydrozoan cnidarians (e.g., *Hydra)* (Chera et al., 2009; Otto and Campbell, 1977) or jellyfish (Lilley et al., 2014). In contrast, most biomedical model organisms (i.e., mammals, nematodes and flies) have fixed, genetically pre-determined body sizes, uncoupling growth from feeding at maturity. In these animals, starvation depletes stored nutrients, causes tissue atrophy (e.g., muscle), and arrests stem cell proliferation arrest in germline or high-turnover tissues, such as the intestinal epithelium (Goodlad and Wright, 1984; McCue, 2010; McLeod et al., 2010). Limited data from animals with lifelong growth leave the cellular, molecular and physiological links between nutrient availability and growth regulation poorly understood, especially beyond embryonic development, regeneration or injury response.

On a cellular level, the nutritional regulation of the cell cycle is well studied in unicellular eukaryotes such as yeast (Breeden and Tsukiyama, 2022; Broach, 2012; Klosinska et al., 2011), green algae (*Chlamydomonas reinhardtii)* (Takeuchi and Benning, 2019), and mammalian cell cultures (e.g., fibroblasts) (Coller et al., 2006; Fujimaki et al., 2019). In these systems, nutrient depletion typically induces cellular quiescence – a reversible cell cycle arrest in G_1_/G_0_ or, more rarely, in G_2_ (Coller et al., 2006; Fujimaki et al., 2019; Sun and Gresham, 2021; Takeuchi and Benning, 2019). In vertebrates and flies, cellular quiescence is predominantly observed in stem cells (e.g., muscle, hematopoietic, neural, intestine) that reactivate upon regeneration, injury or growth factor stimulation. Notably, despite their quiescence, adult mammalian stem cells occasionally divide to support tissue renewal, enabling their identification by long-term label retention (Li and Clevers, 2010; van Velthoven and Rando, 2019).

Nutritionally regulated quiescence has been identified in few cases, such as larval neuroblasts in *Drosophila* (Britton and Edgar, 1998; Otsuki and Brand, 2018), neural progenitors in *Xenopus* (McKeown and Cline, 2019) and germline stem cells in *C. elegans* (Seidel and Kimble, 2011; Seidel and Kimble, 2015), as well as in the adult mouse brain (Otsuki and Brand, 2020). The TOR signalling pathway, largely conserved among eukaryotes, controls nutritional reactivation of the cell cycle in some animals (e.g. *Drosophila*, *Xenopus*) (Chell and Brand, 2010; McKeown and Cline, 2019; Sousa-Nunes et al., 2011) but not others (e.g., *C. elegans*) (Seidel and Kimble, 2015). The predominance of non-nutritional cues in regulating stem cell quiescence and proliferation in animals with fixed body sizes has led to speculations that with the evolution of multicellularity, quiescence became controlled by secreted growth factors rather than nutrients (Daignan-Fornier et al., 2021; Urbán and Cheung, 2021). Since indeterminately growing animals depend on nutrient-regulated growth control, studying animals like sea anemones may be key to understanding the significance and evolution of starvation-induced cellular quiescence.

Although cellular quiescence was discovered decades ago, its metabolic and genetic regulation has only recently begun to be elucidated (Augenlicht and Baserga, 1974; Epifanova and Terskikh, 1969; Owen et al., 1989). In general, quiescence is an actively maintained state characterized by increased chromatin compaction and reduced metabolic, transcriptional and translational activity in both animals and yeast (Cheung and Rando, 2013; de Morree and Rando, 2023; Sun and Gresham, 2021). Work in mammalian cell culture has introduced the concept of ‘quiescence depth’ to explain variations in the degree of quiescence, where cells in deeper quiescence require stronger stimuli, exhibit delayed cell cycle re-entry, and show greater transcriptomic shifts than those in shallow quiescence (Augenlicht and Baserga, 1974; Coller et al., 2006; Fujimaki et al., 2019; Kwon et al., 2017; Laporte et al., 2017; Liu et al., 2024; Marescal and Cheeseman, 2020; Owen et al., 1989). Prolonged deprivation of nutrients or growth factors can lead to loss of proliferative competence and/or to senescence in cultured cells (Fujimaki et al., 2019; Fujimaki and Yao, 2020; Sun and Gresham, 2021). Similar variations in ‘readiness’ to re-enter the cell cycle were found in adult mammalian stem cells, with increased TOR complex 1 activity marking the transition from G_0_ to ‘G_alert_’ or a primed state (van Velthoven and Rando, 2019). Recent studies in fibroblast cell cultures indicate that quiescence depth is modulated by the duration of nutrient or growth factor deprivation and is regulated by autophagy, lysosomal activity and the Retinoblastoma/E2F transcription factor network (Fujimaki et al., 2019; Fujimaki and Yao, 2020; Kwon et al., 2017). Whether the nutritional control of quiescence depth has organismal relevance, particularly in the context of body size plasticity and growth regulation, remains unclear.

On a whole-body level, the relationship between feeding, cell cycle dynamics and body size plasticity has been mainly studied in *Hydra* and planarians (e.g., *Schmidtea mediterranea*) (Figure 1A) (Baguñà et al., 1990; Bosch and David, 1984; Böttger and Alexandrova, 2007; Chera et al., 2009; David and Campbell, 1972; González-Estévez et al., 2012; Newmark and Alvarado, 2000; Otto and Campbell, 1977). In starved *Hydra*, interstitial and epithelial stem cells remain slowly proliferative, with extended S and G_2_ phases enabling rapid cell cycle acceleration upon refeeding (Buzgariu et al., 2014; David and Campbell, 1972; Herrmann and Berking, 1987). Similarly, in planarians, pluripotent neoblasts continue dividing during starvation (Baguñà, 1974; Baguñà, 1976; González-Estévez et al., 2012). In both organisms, starvation-induced whole-body shrinkage results from increased cell loss relative to proliferation, without apparent quiescence in G_1_ (Bosch and David, 1984; González-Estévez et al., 2012).

**Figure 1.**
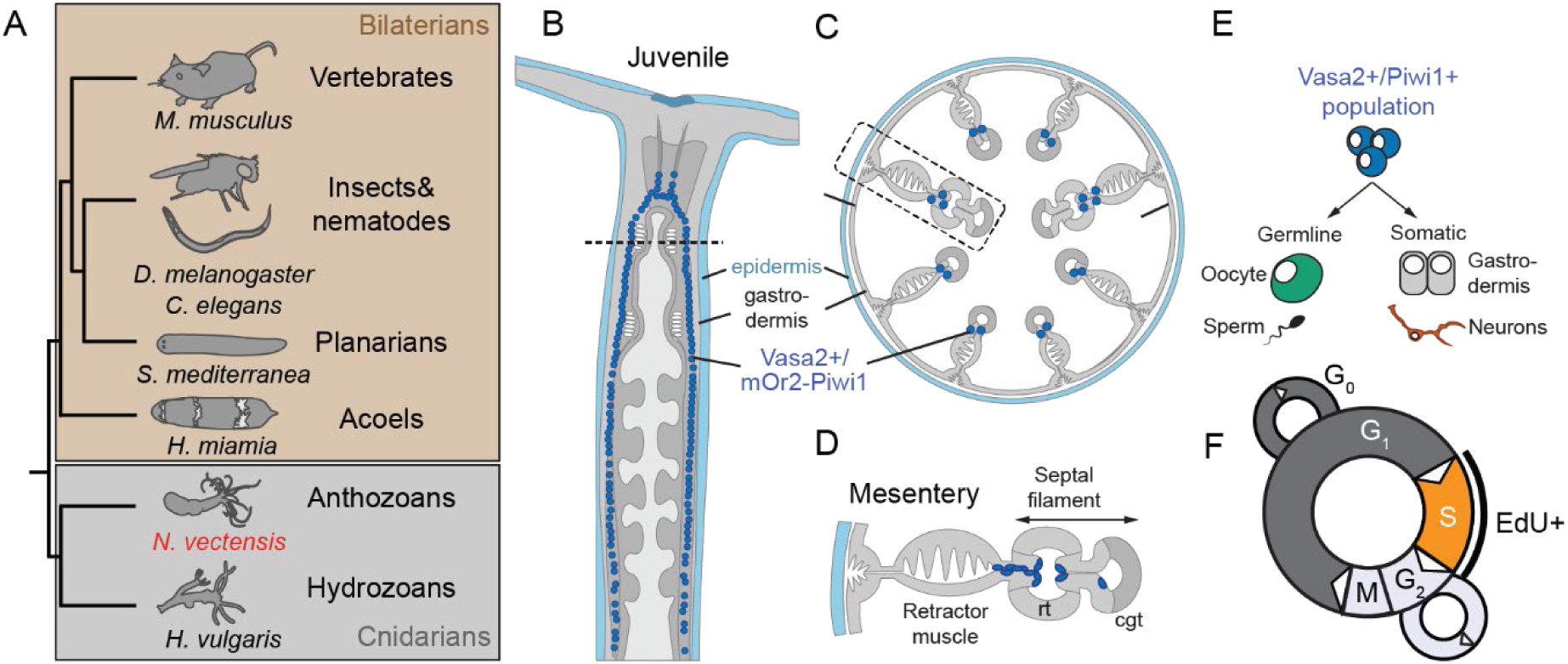
The phylogenetic position of *Nematostella* and localisation of Vasa2+/Piwi1+ cells within the juvenile body plan. (A) Simplified phylogenetic tree highlighting the phylogenetic position of the sea anemone *Nematostella vectensis* and other animal taxa relevant for this study. All animal silhouettes are licensed under CC0 1.0 Universal Public domain and taken from https://www.phylopic.org. (B-D) Schematics showing the localisation of Vasa2+/Piwi1+ cells in a juvenile polyp, depicted in longitudinal (B) or cross-section (C-D). (E) Schematic representation of the multipotent, Vasa2+/Piwi1+ stem/progenitor cell population and a simplified summary of their germinal and somatic progeny. (F) Schematics of cell cycle phases, highlighting the incorporation of EdU during S-phase (black line). Full species names in (A): *Caenorhabditis elegans*, *Drosophila melanogaster, Hofstenia miamia*, *Hydra vulgaris, Mus musculus*, *Nematostella vectensis* and *Schmidtea mediterranea.* cgt: cnidoglandular tract. rt: reticular tract.

The sea anemone *Nematostella vectensis* has emerged as a valuable research organism for studying body plasticity in response to environmental changes, including nutrient availability and temperature (Al-Shaer et al., 2023; Baldassarre et al., 2022; Garschall et al., 2024; Havrilak et al., 2021; Reitzel et al., 2013). A recent study showed that feeding triggers growth and cell proliferation, while starvation induces whole body shrinkage (Garschall et al., 2024). However, the stem or progenitor cell populations driving this plasticity and their responses to feeding or starvation remain unexplored. We therefore studied a recently identified proliferative cell population in *Nematostella* that co-expresses conserved germline and multipotency genes (e.g., *Vasa2, Piwi1*) (Miramón-Puértolas et al., 2024). These Vasa2+/Piwi1+ cells constitute ∼0.05% of all cells in a fed juvenile. They contribute to both germline and somatic cells, including *soxB(2)+* neuronal progenitor cells, and represent a putative multipotent stem cell population, though their homogeneity remains to be investigated (Figure 1B-1E) (Miramón-Puértolas et al., 2024).

Here, we study the response of Vasa2+/Piwi1+ cells to starvation and re-feeding in juveniles, revealing hallmarks of deepening quiescence under prolonged starvation. We further demonstrate that their cell cycle re-entry upon refeeding depends on TOR signalling. Our findings suggest that the nutritional regulation of quiescence depth, previously characterized in unicellular organisms and cell cultures, may be an evolutionary conserved mechanism underlying animal growth plasticity.

## Results

### The cell cycle of Vasa2+/Piwi1+ cells responds dynamically to food availability

Cell proliferation in juvenile *Nematostella* polyps is tightly regulated by food supply. Feeding induces a transient burst of proliferation throughout the polyp, which subsides to low baseline levels within two days (Garschall et al., 2024). To investigate how the Vasa2+/Piwi1+ cell population responds to nutrient availability, we focused on quantifying changes in their relative abundance, proliferation and cell cycle phase distribution over 40 days of starvation. We used a transgenic knock-in line expressing mOr2-Piwi1 protein in Vasa2+/Piwi1+ cells (Miramón-Puértolas et al., 2024) and gated cells with high mOr2-Piwi1 intensities by flow cytometry (FC) to study the responses of Vasa2+/Piwi1+ cells. The proportion of Vasa2+/Piwi1+ cells increased by ∼2.2-fold between T_0_ and 20 days of starvation (Figure 2A; Table S1C) and by ∼6.2-fold between 20 and 40 days of starvation (Figure 2A, 2B and S1; Table S1A-C and S1F).

**Figure 2.**
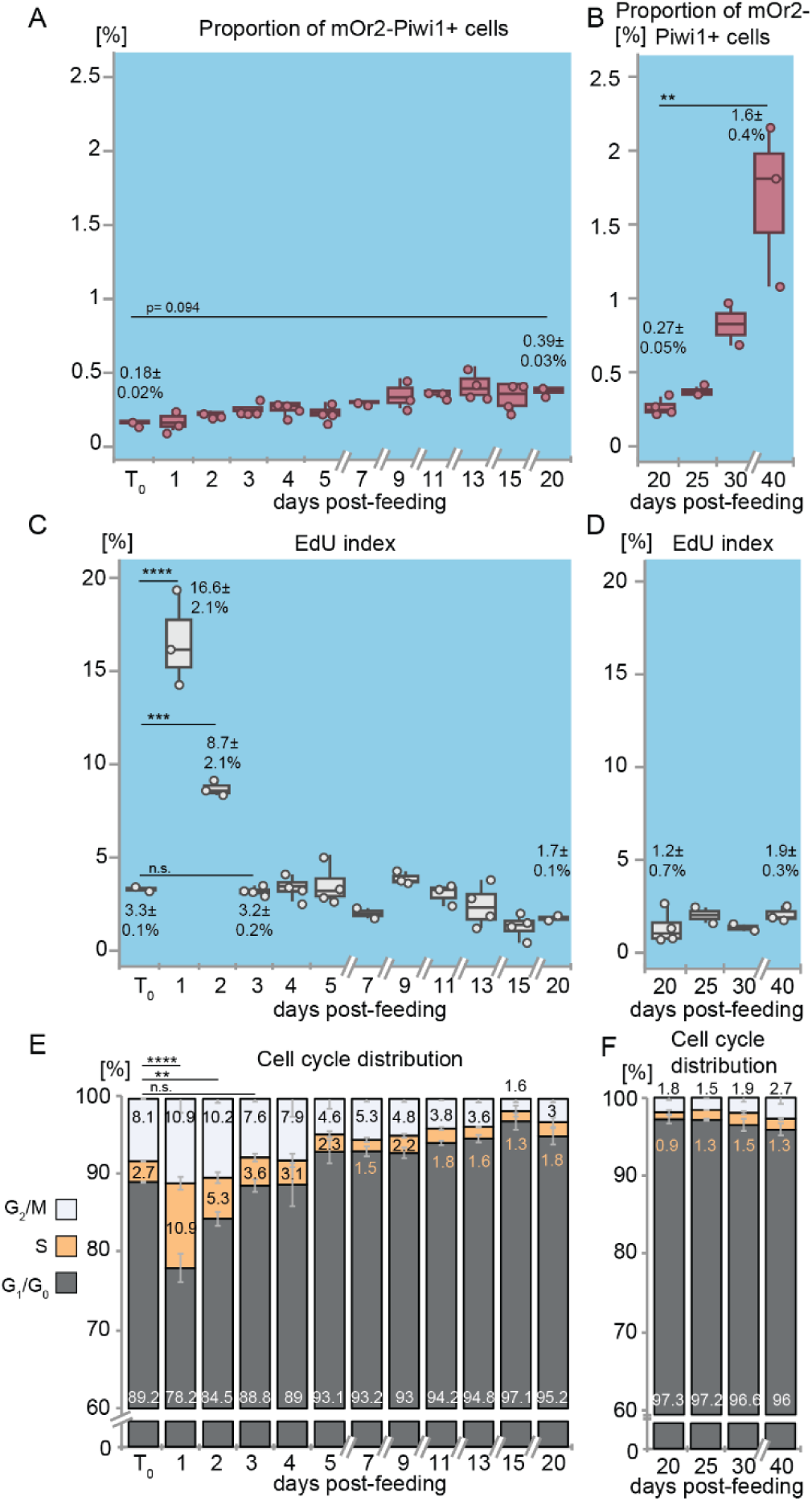
Feeding and starvation affect the proportion and cell cycle activity of Vasa2+/Piwi1+ cells. (A, B) The proportion of mOr2-Piwi1+ cells increases between T_0_ and 20 days (A), and between 20 and 40 days of starvation (B). (C-F) Refeeding at T_0_ triggers a transient peak in cell proliferation as indicated by changes in the proportion of EdU+ cells (EdU index; C, D) and cell cycle phase distributions (E, F) measured by flow cytometry. 5 days starved polyps (T_0_) were refed for 1 hour and sampled at indicated days post-feeding. See ‘Data visualisation’ for definition of box plots and bar plots. Values in A-D represent means ± standard deviations of respective timepoints with dots representing individual samples. Values in E, F represent means. *n*= 2-4 biological replicates per condition, each replicate consisting of a pool of 15 animals. Significance levels after one-way ANOVA with Tukey’s HSD for pairwise comparisons are indicated for adjusted *p* values: \*\**p*<0.01; \*\*\**p*<0.001; \*\*\*\**p*<0.0001; d: day(s); n.s.: non-significant. See also Table S1.

Proliferation rates were assessed using 30-minute incubation pulses of 5-ethynyl-2’- deoxyuridine (EdU), a thymidine analogue incorporating into DNA during S-phase. DNA content measurements allowed quantification of cells in G_1_/G_0_ (2N DNA content), S (between 2N and 4N), or G_2_/M (4N) cell cycle phases (Figure 1F) (Garschall et al., 2024). At T_0_, when juveniles were starved for 5 days, only 3.3±0.1% of Vasa2+/Piwi1+ cells incorporated EdU, indicating low proliferation. A 1-hour refeeding pulse triggered a ∼5-fold increase in the EdU index and a ∼4-fold rise in S-phase cells within 24h. Both values halved by 48h and returned to baseline within three days (Figure 2C and 2E; Table S1A, S1D and S1E). The simultaneous decrease in G_1_/G_0_- and minor increase in G_2_/M-phase cell proportions (Figure 2E) indicate that refeeding prompted cell cycle re-entry from G_1_/G_0_. The sharp, transient surge in the EdU index and S-phase proportion in Vasa2+/Piwi1+ cells further suggests that S-phase re-entry occurs relatively synchronously

Between 3 and 40 days of starvation, the proportion of Vasa2+/Piwi1+ cells in S-phase remained low without showing another transient peak of S-phase progression (EdU: 1.2-3.2%; S: 0.9-3.6%) (Figure 2D and 2F; Table S1A, S1B, S1D, S1E, S1G and S1H). These findings suggest that Vasa2+/Piwi1+ cells continue to proliferate at constant but low rates during prolonged starvation.

### Feeding duration determines proliferative competence

While feeding induced a sharp but transient proliferation burst, it remained unclear whether all or only a subset of Vasa2+/Piwi1+ cells responded to the feeding stimulus. To investigate how starvation and refeeding durations impact their proliferative competence, we assessed their re-entry following different starvation periods. We used continuous EdU incubation over 5-7 days to determine the cumulative EdU index (cEdU), which reflects the proportion of cells that underwent at least one S-phase since T_0_ (i.e., after 5 or 20 days of starvation). We found that regardless of the preceding starvation duration, in the continued absence of food, ∼30-36% of Vasa2+/Piwi1+ cells incorporated EdU over 7 days, indicating that starvation duration has no major effect on spontaneous proliferation rates (Figure 3A and S2; Table S2A and 2L).

**Figure 3.**
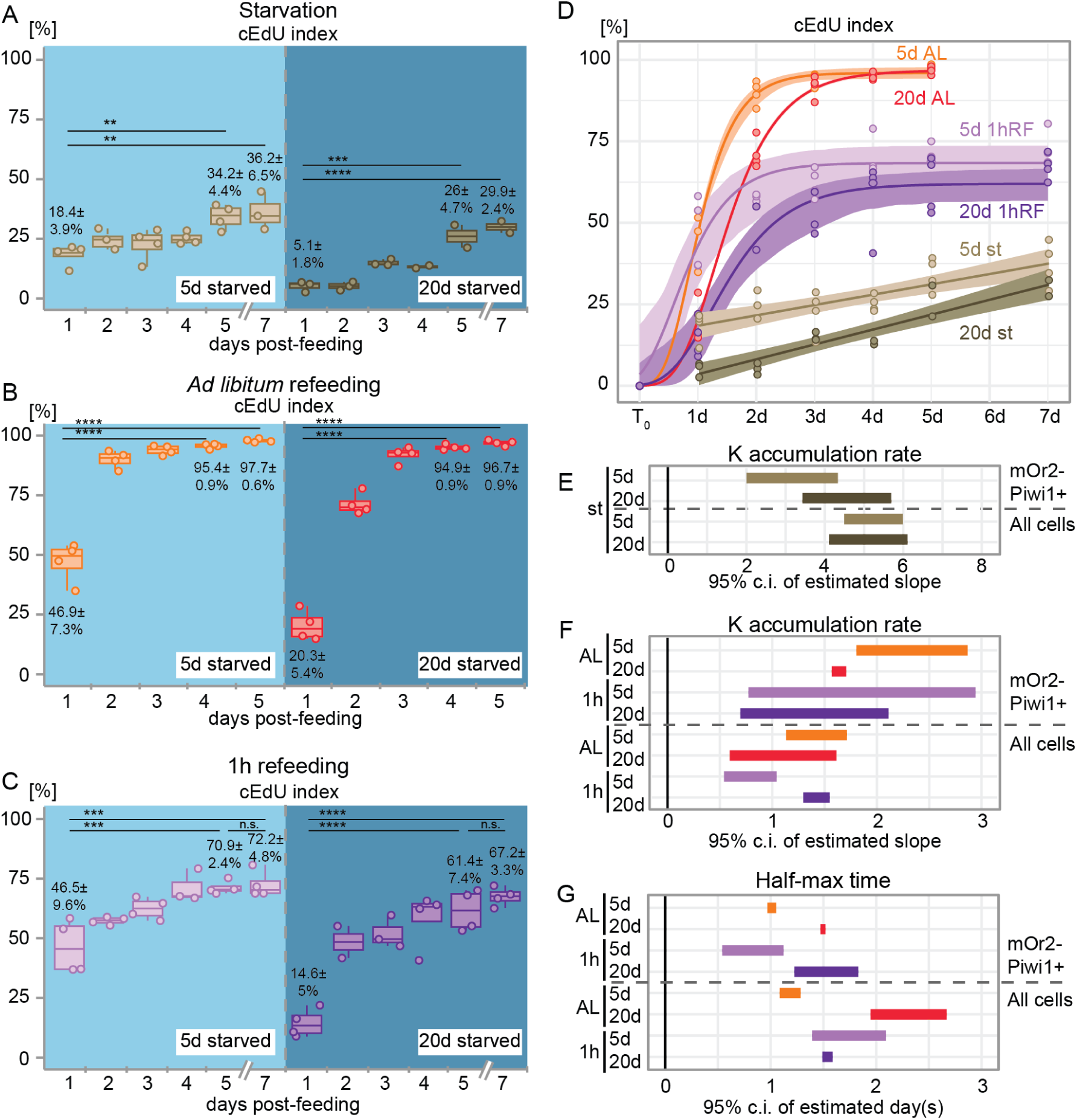
The effect of feeding and starvation on the proliferative competence and onset of cell cycle re-entry in Vasa2+/Piwi1+ cells. (A-C) Temporal changes in the cumulative EdU (cEdU) index under continued starvation (A), *ad libitum* refeeding (B) or following a single, 1-hour refeeding pulse (C) after 5 or 20 days of starvation. Experiments were done using flow cytometry. (D) Dynamics of the cEdU index (A-C) are best explained by linear growth models under continued starvation (st), or by Gompertz growth models after *ad libitum* (AL) or a 1-hour refeeding pulse (1hRF). Dots represent same replicate sample values as in (A-C). (E, F) Estimating the effect of starvation (E), *ad libitum* and 1-hour refeeding (F) on accumulation rate of cEdU+ cells (K) was done by comparing the 95% confidence intervals between 5 and 20 days of starvation. K in (E) derived from linear models (D), and (F) derived from Gompertz growth models (D). (G) Estimating the effect of *ad libitum* and 1-hour refeeding on the half-max time, when the cEdU index reaches 50% of its maximum, was done by comparing the 95% confidence intervals between 5 and 20 days of starvation. Half-max time derived from Gompertz growth models (D). ‘All cells’ refers to all cell cycle-gated cells (incl. mOr2-Piwi1+; Figure S2). *n*=2-4 biological replicates per condition (15 individuals per replicate). Coloured lines in D represent the model curve or line for each condition with overlays depicting 95% confidence intervals. See ‘Data visualisation’ for definition of box plots. Dots represent individual values. Index values represent means ± standard deviations of respective timepoints. Pairwise comparisons after one-way ANOVA were calculated using Tukey’s HSD and *p* values adjusted at significance codes: \*\**p<*0.01; \*\*\**p<*0.001; \*\*\*\**p<*0.0001. d: day(s), n.s.: non-significant. See also Figure S3 and S4, and Table S2, S3 and S4.

After 5 days *ad libitum* (AL) refeeding, ∼96-97% of Vasa2+/Piwi1+ cells incorporated EdU, independent of prior starvation duration (Figure 3B; Table S2D and S2K). This contrasts with the ∼13-25% lower cEdU index observed among all polyp body cells, which includes terminally differentiated cells that cannot re-enter the cell cycle (Figure S3B; Table S3D and S3K). Following a single 1-hour refeeding pulse, the cEdU index measured after 5 or 7 days was well above starvation baseline but significantly lower than during AL refeeding (Figure 3A-3C; Table S2A, S2D, S2G and S2K). Notably, starvation duration had no significant effect on the proportion of Vasa2+/Piwi1+ cells capable of re-entering the cell cycle, but a single refeeding stimulus resulted in a lower proliferative competence than AL refeeding (Figure 3B and 3C).

### Feeding stimulus and starvation history affect the dynamics of cell cycle re-entry

We then explored how starvation history affects the dynamics of EdU+ cell accumulation by comparing 24h-cEdU values across feeding regimes and starvation durations. Notably, under the same feeding regime (starvation, AL or 1-hour), the fraction of Vasa2+/Piwi1+ cells incorporating EdU within 24h was consistently 2-3 times higher after 5 than after 20 days of starvation (Figure 3A-3C; Table S2A, S2D, S2G and 2J). However, refeeding either AL or 1-hour after the same starvation duration (5 or 20 days) consistently yielded similar cEdU values within 24h of refeeding (Figure 3B and 3C; Table S2D, 2G and 2J). These results suggest that starvation history but not feeding regime affects the proportion of cells re-entering the cycle during the first 24h after refeeding.

To determine whether these differences arise from changes in the rate or onset of proliferation, we estimated cEdU index dynamics by applying linear regression or growth models to the data. During continuous starvation, a single-phase linear regression model proved a good fit to the datapoints between 1 and 7 days of EdU incubation (5d: R^2^=0.6063; 20d: R^2^=0.8543, Table S4A). We observed that independent of the starvation history, the cEdU index linearly increased between 1 and 7 days without apparent stagnation (5d_Δd7/1_: ∼2-fold; 20d_Δd7/1_: ∼6-fold). Therefore, all Vasa2+/Piwi1+ cells may eventually proliferate even under prolonged starvation. Extrapolation predicted that after 5 and 20 days of starvation, all Vasa2+/Piwi1+ cells would have entered S-phase after 31.5 and 21.9 days, respectively. Notably, the cEdU accumulation rate (K) was significantly higher (i.e., steeper slope) after 20 than 5 days of starvation, with minimal overlap in their 95% confidence intervals (Figure 3D and 3E; Table S4A). Interestingly, K_20d_ of Vasa2+/Piwi1+ cells resembled values from all polyp cells as their 95% confidence intervals largely overlapped (Figure 3E and S3D; Table S4A). These findings suggest that EdU+ cell accumulation in Vasa2+/Piwi1+ cells is unusually slow after short starvation but normalizes following prolonged starvation to a level generally observed among all proliferating cells. Differences in accumulation rates between short and long starvation may reflect changes in the cell division symmetry among Vasa2+/Piwi1+ cells (see Discussion).

After AL refeeding, a Gompertz Growth model (Gompertz, 1825; Winsor, 1932) proved an excellent fit to describe the dynamics of measured cEdU indices (5d: R^2^=0.999; 20d: R^2^=0.998; Table S4B; Figure S4A). From this model, we estimated the accumulation rate K and half-max time t50, which represents the timepoint at which the cEdU index reaches 50% of its maximum (Figure S4C). Following AL refeeding, K_5d_ was significantly higher than K_20d_, with non-overlapping 95% confidence intervals (Figure 3F; Table S4C), while t50 differed by about half a day (Figure 3G; Table S4C). Similarly, among all cells (Figure S4B), AL refeeding after long starvation led to reduced cEdU accumulation rates and delayed t50 by about a full day (Figure 3F, 3G and S3D; Table S4C). Together, these results indicate that longer starvation slows and delays EdU+ cell accumulation upon AL refeeding.

To assess whether this pattern persisted following a 1-hour refeeding stimulus, we again fitted a Gompertz Growth model (5d R^2^=0.9685; 20d R^2^=0.9796; Table S4B; Figure S4A and S4B). While K values did not differ significantly, t50 was significantly delayed after 20 days of starvation, mirroring AL refeeding results (Figure 3F and 3G; Table S4C). Together, these findings suggest that starvation history affected the onset but not the rate of accumulation under any refeeding regime.

### Starvation length affects onset of cell cycle re-entry

To assess whether lower K and t50 values after prolonged starvation reflect delays in cell cycle re-entry, we analysed the 30min-EdU ‘snapshot’ index at high temporal resolution following a single 1-hour refeeding stimulus. To control for potential circadian effects, we found that sampling in the morning or evening did not significantly affect EdU indices, the proportion of phospho-histone H3+ cells (Chen et al., 2020; Hendzel et al., 1997) (Figure S5A) or cell cycle distributions of Vasa2+/Piwi1+ cells from polyps starved for 5 or 20 days (Figure S5B-S5D and S6).

Following 5 days of starvation, the EdU index increased significantly within 6 hours of refeeding, whereas after 20 days of starvation, a significant increase was only observed at 18 hours and onwards (Figure 4A; Table S5A, S5C and S5F). This 12-15-hour delay in S-phase re-entry was further confirmed by flow cytometry-based DNA content analysis (Figure 4B; Table S5A, S5D and S5G). While the EdU index at T_0_ was not significantly different (Figure 4A; Table S5A and S5B), the proportions of S and G_2_/M phase cells were markedly lower after prolonged starvation (Figure 4B; Table S5A and S5B). After short-term starvation, the proportion of S-phase cells significantly increased already within 3 hours of re-feeding, whereas after 20 days of starvation, a significant increase was only detected only after 21 hours (Figure 4B; Table S5A, S5D and S5G). These findings strongly suggest that prolonged starvation delays S-phase re-entry in Vasa2+/Piwi1+ cells.

**Figure 4.**
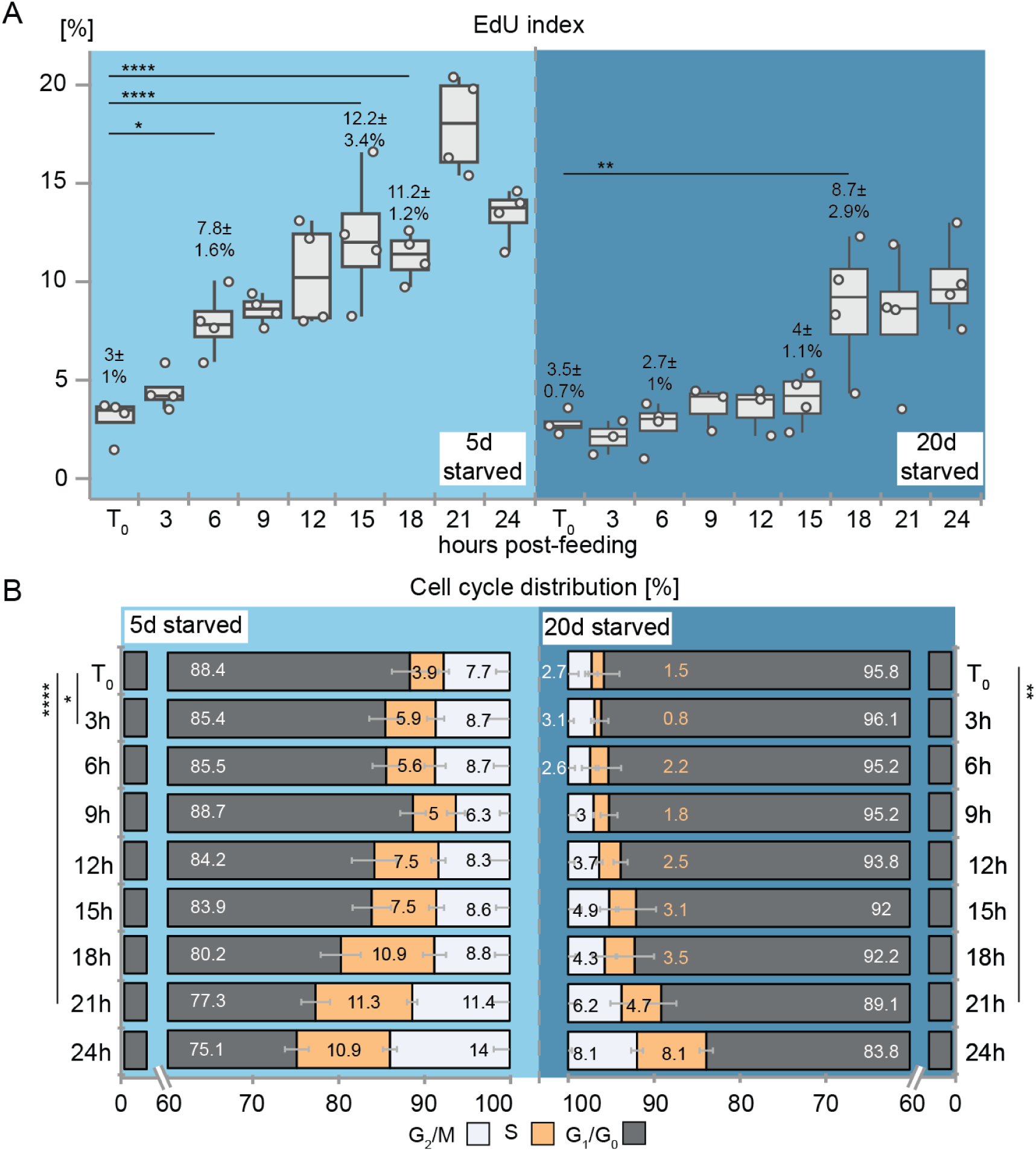
Prolonged starvation leads to a delay in cell cycle re-entry in Vasa2+/Piwi1+ cells. (A, B) Proportion of EdU+ cells (EdU index; A) and cell cycle phase distributions (B) over 24 hours after refeeding polyps starved 5 or 20 days measured by flow cytometry. The earliest significant changes in the EdU index (A) and S-phase proportion (B) to T_0_ are indicated, highlighting the delayed cell cycle re-entry after 20 days of starvation. Note that T_0_ between 5 and 20 days exhibits major differences in the proportions of S and G_2_/M phase cells. Starved polyps (T_0_) were refed for 1 hour. For box plots and bar plot definitions, see ‘Data visualisation’. Values in A represent means ± standard deviations of respective timepoints with dots indicating individual samples. Values in B represent means. *n*= 2-4 biological replicates per condition, with 15 polyps per replicate. Significance levels after one-way ANOVA with Tukey’s HSD for pairwise comparisons are indicated for adjusted *p* values: \**p<*0.05, \*\**p<*0.01; \*\*\*\**p<*0.0001. d: day(s), n.s.: non-significant. See also Figure S5 and Table S5.

We also quantified the proportion of mitotic, pH3+ cells to determine if the delay in re-entry was linked to the reduced proportion of G_2_/M phase cells. No significant changes in pH3 index were observed in either starvation condition within the first 15 hours post-feeding, suggesting that synchronous cell cycle re-entry is not driven by G_2_/M-phase cells but rather by a relatively synchronous re-entry from G_1_/G_0_ (Figure S5E; Table S5A, S5E and S5H).

### Starvation length affects feeding-dependent TOR signalling in Vasa2+/Piwi1+ cycling cells

To investigate the molecular mechanisms of delayed cell cycle re-entry after prolonged starvation, we examined whether the target of rapamycin (TOR) signalling activity correlate with these differences. We used the phosphorylation of the ribosomal protein S6 (pRPS6) by S6 kinase as a readout, a well-established marker of active TOR signalling in yeast (Yerlikaya et al., 2016), sea anemones (Garschall et al., 2024; Ikmi et al., 2020; Voss et al., 2023) and bilaterians (Magnuson et al., 2012). Using flow cytometry (Figure S7), we quantified the proportion of EdU+ and pRPS6+ cells (pRPS6 index) over time to assess TOR activity in relation to S-phase re-entry after short- and long-term starvation. To rule out circadian effects, we confirmed that cell proliferation and pRPS6 index levels did not significantly differ between Vasa2+/Piwi1+ cells from 5- or 20-day starved polyps sampled 9 hours apart (Figure S8A-S8C; Table S6A and S6B). At T_0_, the pRPS6 indices between short and long starvation were not significantly different (Figure S8A; Table S6A and S6B).

Following 1-hour refeeding, the pRPS6 indices showed considerable variation at individual time points, but overall, the dynamics between short- and long-term starvation differ (Figure 5A, C; Table S6C). After short starvation, pRPS6 indices increased within hours and remained high (20-30%) up to 48 hours post-refeeding. In contrast, after 20 days of starvation, pRPS6 indices initially declined to <5% over 9 hours before reaching 20-30% only at 48h (Figure 5A and 5C; Table S6C). The EdU index dynamics confirmed the previously observed delay in S-phase re-entry after long starvation, with a slightly earlier increase (at 15 hours, Figure 5B and 5D; Table S6C) compared to the identified previously 18-hour timepoint (Figure 4A). The temporal changes in pRPS6 levels closely mirrored those of the EdU index. Pearson correlation analysis showed a positive correlation between pRPS6 and EdU indices over 72 hours after short starvation (Figure 5A and 5B; Table S6C and S6D), but only during the first 30 hours after long starvation (Figure 5C and, 5D; Table S6C and, S6E-S6G). These findings suggest that TOR signalling activity is closely associated with S-phase re-entry, particularly early after re-feeding, raising the question of whether TOR acts in parallel or upstream of S-phase re-entry and whether it is predominantly active in proliferating cells.

**Figure 5.**
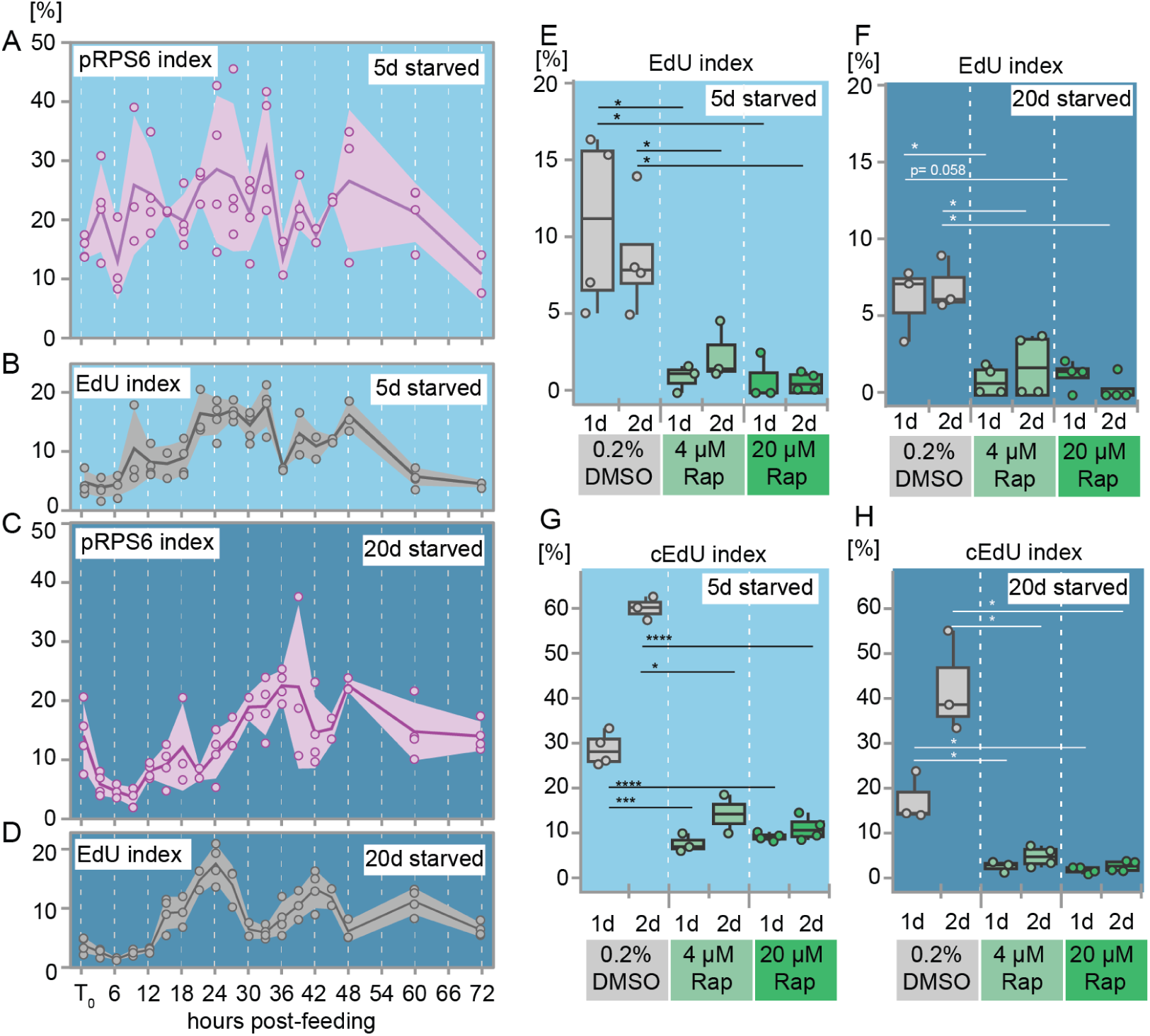
Dynamics and functional role of TOR signaling during cell cycle re-entry in Vasa2+/Piwi1+ cells. (A-D) Changes in the proportion of pRPS6+ (pRPS6 index; A, C) and EdU+ cells (EdU index; B, D) over 72 hours after refeeding of 5- (A, B) or 20-days starved polyps (C, D). Experiments were done using flow cytometry. Note similarities in the pRPS6 and EdU index dynamics within the first 24h after feeding. Coloured lines indicate mean per condition and band overlays represent 95% confidence intervals. *n*= 2-4 biological replicates per condition, with 15 individuals per replicate. Dots represent individual values. (E-H) The TOR inhibitor Rapamycin (’Rap’) decreases the proportion of EdU+ (EdU index; E, F) and cumulative EdU+ cells (cEdU index; G, H) within 1 and 2 days after refeeding compared to DMSO controls. For box plots, see ‘Data visualisation’. Dots represent individual values. *n*= 2-4 biological replicates per condition, with 15 individuals per replicate. Significance levels for Student’s t-test are indicated for *p* values: \**p*<0.05, \*\*\**p*<0.001, \*\*\**p*<0.0001. d: day(s). See also Figure S8 and S10, and Table S6 and S7.

Confocal imaging confirmed that pRPS6 protein is present in both S-phase (EdU+) and M-phase (metaphase plate) Vasa2+/Piwi1+ cells (Figure S8D-S8K). To determine whether pRPS6+ were enriched in specific cell cycle phases, we analysed cell cycle distributions within pRPS6+ and pRPS6– fractions (Figure S9A and S9B; Table S6H). Calculating the log_2_FC of their ratio (pRPS6+/pRPS6–), we found across nearly all timepoints that S- and G_2_/M-phase cells were overrepresented (log_2_FC>0), while G_1_/G_0_ cells were underrepresented (Figure S9C and S9D; Table S6I). These results indicate that proliferative Vasa2+/Piwi1+ cells predominantly exhibit active TOR signalling, regardless of starvation duration.

### TOR signalling is required for feeding-dependent cell proliferation in Vasa2+/Piwi1+ cells

To test whether TOR signalling is functionally required for refeeding-induced cell cycle re-entry, we treated polyps with two independent TORC1 inhibitors: the allosteric inhibitor Rapamycin (’Rap’) and the ATP-competitive inhibitor AZD-8055 (’AZD’). Western Blot analysis confirmed that both inhibitors strongly suppressed pRPS6 levels at two concentrations, with Rapamycin being more effective (Figure S10A-S10D; Table S7A-S7C and S8).

EdU incorporation assays showed that both TOR inhibitors significantly reduced the EdU index and S-phase re-entry in Vasa2+/Piwi1+ cells after refeeding, irrespective of starvation length (Figure 5E, 5F, S1 and S10E; Table S7D, S7E, S7G and S7H). Flow cytometry analysis further revealed that TOR inhibition reduced S-phase and G_2_/M-phase cell proportions while increasing G_1_/G_0_ proportions (Figure S10F and S10G; Table S7D, S7F, S7G and S7I). Notably, Rap and AZD had a stronger effect on the EdU index than on the S-phase proportion determined by DNA content. To test whether a subset of cells was arrested in S-phase or continued proliferating under Rapamycin treatment, we assessed the cEdU index over 24 hours of continuous EdU incubation. We observed a marked reduction in the cEdU index under Rap, with a stronger effect after long starvation (Figure 5G, 5H and S2; Table S7J and S7K). Notably, the cEdU index at 1- and 2-days post-refeeding (Figure 5G and 5H; Table S7J) remained higher than the corresponding EdU index, regardless of starvation length (Figure 5E and 5F; Table S7D). Our findings show that refeeding-induced cell cycle re-entry in Vasa2+/Piwi1+ cells depends on TOR signalling. However, a small subset of Rapamycin-insensitive Vasa2+/Piwi1+ cells continues to cycle after refeeding, with its proportion decreasing under prolonged starvation.

## Discussion

In the sea anemone *Nematostella vectensis*, body size and growth rates are tightly linked to nutrient availability, characteristic of animals with lifelong growth. Starvation leads to considerable cell loss and body shrinkage, while refeeding triggers cell proliferation and growth (Garschall et al., 2024). To understand how food availability regulates growth at the cellular level, we studied the nutritional regulation of Vasa2+/Piwi1+ stem/progenitor cells (Miramón-Puértolas et al., 2024). Their proportion increased ∼9.4-fold over 40 days of starvation, despite a >50% reduction in polyp size and total cell numbers (Garschall et al., 2024). This accumulation may result from either protection from cell loss or expansion by continued proliferation.

Cell proliferation among Vasa2+/Piwi1+ cells was highly dependent on food availability. Refeeding after 5 days starvation induced a proliferation surge within 24 hours, indicated by a rapid increase in the EdU index and relative proportion of S- and G_2_/M phases. This burst was followed by a decline in the EdU index and S-phase proportion, with a complementary increase of G_1_/G_0_-phase cells within three days. These dynamics suggest that refeeding triggers a relatively synchronous re-entry into S-phase from a quiescent G_1_/G_0_ state (Figure 6). Beyond 3 days of starvation, the proportion of S-phase/EdU+ cells remained stable at approximately 1-3.5%, indicating spontaneous, rather than synchronized, cell divisions. This suggests that daily feeding induces a circadian rhythm in cell cycle progression, which is lost during starvation, similar to observations in *Hydra* stem cells (Buzgariu et al., 2014). Together, our observations indicate that starvation and refeeding induce quiescence and cell cycle re-entry, respectively, in Vasa2+/Piwi1+ cells (Figure 6).

**Figure 6.**
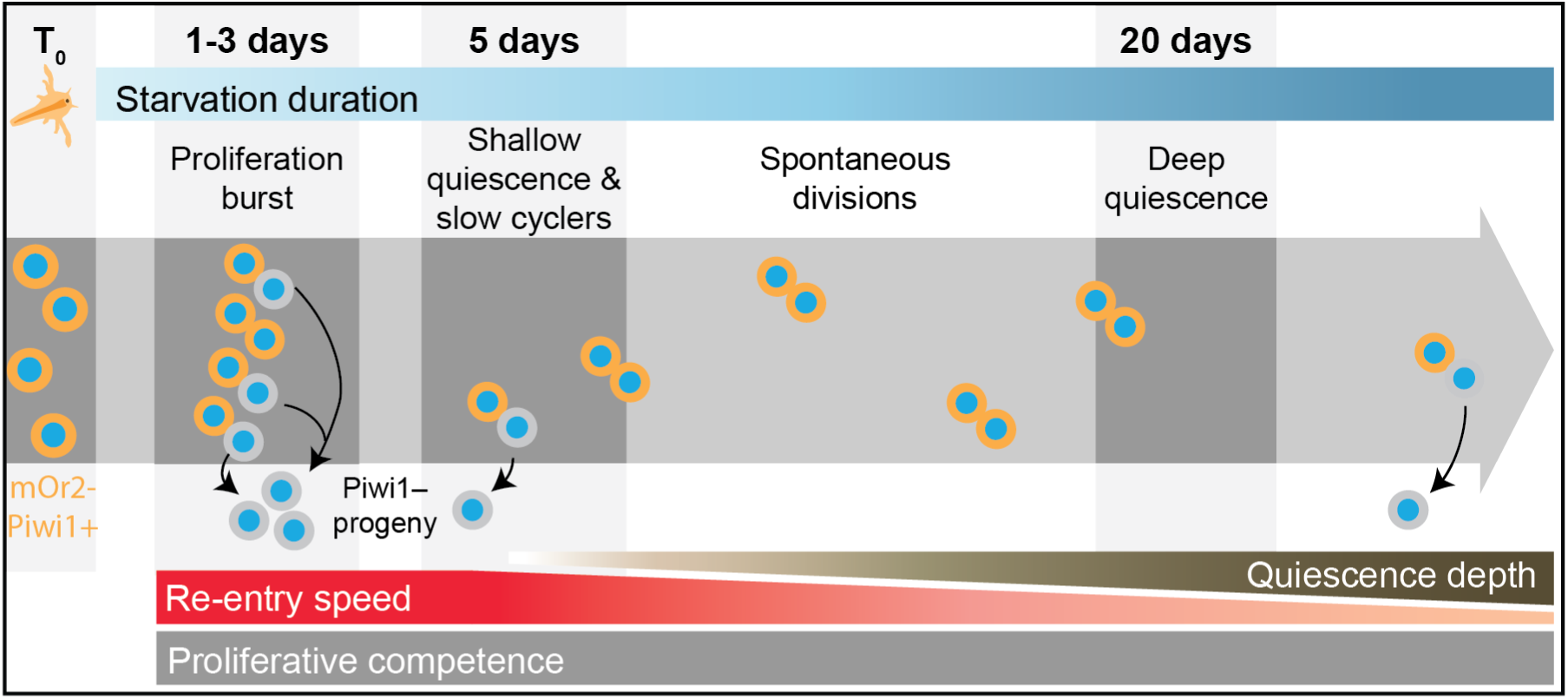
Working model of the nutritional regulation of Vasa2+/Piwi1+ cell proliferation and quiescence. Our work indicates that feeding triggers a burst of cell proliferation in Vasa2+/Piwi1+ cells, while starvation induces cellular quiescence. Prolonged starvation deepens quiescence, characterised by a delayed cell cycle re-entry and TOR signalling activity increase upon refeeding. After short-term starvation (5 days), a subset continues to divide slowly and asymmetrically. Under continued starvation, some cells spontaneously exit quiescence and divide symmetrically. Even after 20 days of starvation, there was no detectable decline in the competence of Vasa2+/Piwi1+ cells to re-enter the cell cycle upon refeeding.

While the EdU index among Vasa2+/Piwi1+ cells remained stable beyond 3 days of starvation, changes in cEdU dynamics and cell cycle phase distributions suggest altered cell cycle lengths and division patterns between 5 and 20 days of starvation. Notably, their 7-day cEdU+ rate increased between 5 and 20 days of starvation, aligning with the rate observed among all cells. This shift may reflect changes from predominantly asymmetric division after short starvation to symmetric divisions as quiescence deepens. A previous study has shown that Vasa2+/Piwi1+ cells produce progeny with depleted mOr2-Piwi1 protein due to asymmetric cell division (Miramón-Puértolas et al., 2024). Flow cytometry therefore excludes those cells from the Vasa2+/Piwi1+ cell pool, which may explain the lower cEdU+ index at 5 days. A shift towards symmetric divisions, where both daughter cells keep high mOr2-Piwi1 protein levels, may explain the relative increase in the cEdU index after 20 days of starvation (Figure 6).

Between 5 and 20 days of starvation, both the cEdU index and G_2_/M proportion declined with a corresponding increase in G_1_/G_0_ cells. In apparent contradiction, the proportion of S-phase cells, as determined by the EdU ‘snapshot’ index, remained stable beyond 3 days of starvation. This discrepancy likely reflects a faster cell cycle progression at 5 days compared to 20 days, allowing more cells to pass through S-phase within 24 hours after short starvation. It also indicates that the subset of slowly proliferating Vasa2+/Piwi1+ cells overall decreases between 5 and 20 days of starvation. Under prolonged starvation, proliferative activity is persistently low but not restricted to a small subset of cells. Together, our observations indicate that quiescent Vasa2+/Piwi1+ cells spontaneously exit quiescence at low rates to potentially support tissue renewal (Figure 6), as seen in mammalian intestinal or hematopoietic stem cells (Li and Clevers, 2010; van Velthoven and Rando, 2019).

A hallmark of quiescent cells is their ability to re-enter the cell cycle in response to a specific stimulus. After 5 or 20 days of starvation, approximately 90-97% of Vasa2+/Piwi1+ cells resided in G_1_/G_0_. Within five days of *ad libitum* refeeding, almost all cells (∼96-97%) re-entered the cell cycle from G_1_/G_0_ (Figure 6). The remaining 3-4% cells may be terminally differentiated, senescent, or may need additional stimuli (e.g., injury, growth factors) to exit quiescence.

A comparison between feeding regimes revealed that a single 1-hour refeeding pulse induced a proliferative response equivalent to 24 hours of *ad libitum* refeeding. However, polyps refed *ad libitum* exhibited an increased rate of EdU+ cell accumulation over the following days. This suggests that feeding frequency, rather than the strength of the initial feeding stimulus, determines the proliferative competence of Vasa2+/Piwi1+.

In contrast to yeast and mammalian cell culture systems, where prolonged stimulus deprivation deepens quiescence, starvation in *Nematostella* did not reduce the competence of Vasa2+/Piwi1+ to re-enter the cell cycle (Fujimaki et al., 2019; Kwon et al., 2017; Liu et al., 2024; Su et al., 1996). Even after 20 days of starvation, nearly all Vasa2+/Piwi1+ cells kept their full proliferative potential. Remarkably, however, refeeding after 20 days of starvation significantly delayed the onset of TOR signalling activation (i.e., RPS6 phosphorylation) and S-phase re-entry by 12-15 hours compared to refeeding after 5 days of starvation. This delay explains the differences in cEdU index at 1 day post-refeeding between 5- and 20-day starved animals found at *ad libitum* and 1-hour refeeding conditions. Altogether, we conclude that deepening quiescence in *Nematostella* Vasa2+/Piwi1+ cells is marked by delays in S-phase re-entry and TOR signalling (Figure 6).

The simultaneous increase in RPS6 phosphorylation and S-phase re-entry raised the question if TOR signalling activation is required for or acts in parallel to quiescence exit. Inhibition of TOR complex 1 demonstrated that TOR signalling is necessary for feeding-induced cell cycle re-entry after short and long starvation. However, a subset of Vasa2+/Piwi1+ cells may remain Rapamycin-insensitive after short starvation. Future studies will determine if Rapamycin insensitivity resulted from incomplete pathway inhibition or reflected an inherent property of a subset of slowly cycling cells.

In sea anemones, *Hydra* and planarians, starvation and refeeding trigger body shrinkage or growth through cell proliferation or cell loss (Bode et al., 1973; Chera et al., 2009; Garschall et al., 2024; González-Estévez et al., 2012; Otto and Campbell, 1977). Similar to *Hydra* interstitial stem cells and planarian neoblasts, *Nematostella* Vasa2+/Piwi1+ cells share some key stem cell features, including a high nucleus-to-cytoplasm ratio and the expression of conserved germline multipotency program genes, such as *piwi* or *vasa* genes (Juliano et al., 2010; Miramón-Puértolas et al., 2024). However, *Nematostella* Vasa2+/Piwi1+ cells are much scarcer than planarian neoblast (∼30-40% of all cells) (Baguñà and Romero, 1981) or *Hydra* interstitial stem cells (∼15-20% of all cells) (Bode et al., 1973). Also, unlike neoblasts and stem cells in *Hydra or Clytia, Nematostella* Vasa2+/Piwi1+ accumulate during starvation and exhibit low baseline proliferation during long-term starvation (Baguñà and Romero, 1981; Bode et al., 1973; Chari et al., 2021; González-Estévez et al., 2012). Neoblasts and *Hydra* stem cells, in contrast, maintain or even increase mitotic activity during prolonged starvation (Baguñà, 1974; Baguñà, 1976; Bosch and David, 1984; Buzgariu et al., 2014; David and Campbell, 1972; Eisenhoffer et al., 2008; González-Estévez et al., 2012; Otto and Campbell, 1977). In *Hydra*, the proportion of epithelial stem cells entering S-phase within 48 hours is about 8-fold higher than in *Nematostella* Vasa2+/Piwi1+ cells after 20 days of starvation (∼40% vs. ∼5%) (Bosch and David, 1984). *Hydra* stem cells also exhibit an extremely short or absent G_1_/G_0_ phase, while their G_2_ phase extends without arresting under starvation (Bosch and David, 1984; Buzgariu et al., 2014; Eisenhoffer et al., 2008; Otto and Campbell, 1977). Together, these differences further highlight the tight nutritional control of quiescence and cell cycle re-entry in *Nematostella* Vasa2+/Piwi1+ cells, resembling the ancestral, nutrient-responsive cell cycle dynamics in yeast or mammalian fibroblast cells (Epifanova and Terskikh, 1969; Sun and Gresham, 2021).

Research on quiescence depth as a result of nutrient withdrawal has so far only been observed in yeast and mammalian cell culture. Our findings suggest that the nutritional control of proliferation and quiescence depth may have persisted in some animals, such as sea anemones, to maintain lifelong growth plasticity. It remains to be determined whether the molecular mechanisms regulating quiescence depth in cultured mammalian cells, such as the Retinoblastoma/E2F protein network (Kwon et al., 2017) or lysosomal activity (Fujimaki et al., 2019), are conserved in sea anemones. Our work helps understanding the mechanisms behind environmentally controlled body plasticity and lays the foundation for exploring the metabolic, transcriptomic and epigenetic changes during the reversible transition between proliferation and quiescence.

## Resource availability

### Lead contact

Further information and requests for resources and reagents should be directed to and will be fulfilled by the Lead Contact, Patrick Steinmetz.

### Materials availability

This study did not generate new unique reagents.

### Data and code availability

The flow cytometry files (.fcs) generated for this project have been deposited at the Figshare repository as https://figshare.com/projects/Flow_cytometry_files_Starvation_duration_regulates_cellular_quiescence_depth_and_cell_cycle_re-entry_in_a_sea_anemone/234587, and are publicly available as of the date of Publication. Western blot membranes raw imaging data are available at the Mendeley Data: https://data.mendeley.com/preview/82g2scj6c7?a=16e5b043-9ff3-4f9b-97bc-7f0677839a7a, and are publicly available as of the date of Publication. This paper does not report original code. Any additional information required to reanalyze the data reported in this paper is available from the lead contact upon request.

## Acknowledgements

We thank Alena Pieters and Jens van Bakel for experimental support, all past and present members of the Steinmetz lab, especially Paula Miramón-Puértolas, for support and discussions, and Eilen Myrvold, Brandon Mellin and Lavina Jubek for taking excellent care of the Michael Sars Centre *Nematostella* culture. The flow cytometry was performed at the Flow & Mass Cytometry Core Facility, Department of Clinical Science, University of Bergen. P.R.H.S., K.G. and E.P.-C. received funding from Norges Forskningsråd (234817) to the Michael Sars Centre. P.R.H.S., K.G. received funding from Norges Forskningsråd (335230). E.P.-C. was funded by an EMBO postdoctoral fellowship (ALTF 406-2021). This work was supported by the Michael Sars Centre core budget from the University of Bergen.

## Author contributions

Conceptualization, E.P.-C., K.G. and P.R.H.S.; Methodology, E.P.-C. and P.R.H.S.; Validation, E.P.-C.; Investigation, E.P.-C. and K.G.; Formal analysis, E.P.-C.; Data curation, E.P.-C.; Visualization, E.P.-C. and P.R.H.S.; Writing - Original Draft, E.P.-C. and P.R.H.S.; Writing – Review & Editing, E.P.-C., K.G. and P.R.H.S.; Resources, P.R.H.S.; Funding Acquisition, E.P.-C. and P.R.H.S.; Supervision, P.R.H.S.

## Declaration of generative AI and AI-assisted technologies in the writing process

During the preparation of this work, the authors used ChatGPT (OpenAI, https://chat.openai.com) to draft code for data visualization and analysis, and for editing and polishing of the manuscript. After using this tool, the authors reviewed and edited the content as needed and take full responsibility for the content of the publication.

## Supplemental Information titles and legends

### Supplemental information

Figures S1-S10.

Table S1. Excel file related to Figure 2.

Table S2. Excel file related to Figure 3 and S3.

Table S3. Excel file related to Figure S3.

Table S4. Excel file related to Figure 3, S3 and S4.

Table S5. Excel file related to Figure 4 and S5.

Table S6. Excel file related to Figure 5, S8 and S9.

Table S7. Excel file related to Figure 5 and S10.

Table S8. Excel file related to Figure S10.

## STAR Methods

### Key resources table

| REAGENT or RESOURCE | SOURCE | IDENTIFIER |
| --- | --- | --- |
| Antibodies |  |  |
| Rabbit anti-DsRed | Takara Bio<br>Clontech | Cat# 632496 |
| Mouse anti-mCherry | Takara/AH<br>Diagnostics | Cat# 632543-CLO |
| Rabbit anti-p-RPS6 (Ser235/236) | Cell Signaling<br>Technology | Cat# 4858 |
| Rabbit anti-pH3 (Ser10) | Cell Signaling<br>Technology | Cat# 9701 |
| FluoTag®-X4 anti-GFP | NanoTag<br>Biotechnologies | Cat# N0304 |
| Rabbit anti-Actin | Sigma-Aldrich | Cat# A5060 |
| Secondary goat anti-Rabbit IgG (H+L)<br>coupled with Alexa Fluor 568 | LifeTech | Cat# A11011 |
| Secondary goat anti-Rabbit IgG (H+L)<br>coupled with Alexa Fluor 488 | LifeTech | Cat# A-11008 |
| Secondary goat-anti-Mouse Rabbit IgG<br>(H+L) coupled with Alexa Fluor 568 | LifeTech | Cat# A11011 |
| Secondary goat anti-Rabbit IgG (H+L)<br>coupled with HRP | Abcam | Cat# Ab9705 |
| Chemicals, peptides, and recombinant proteins |  |  |
| FxCycle™ Violet | ThermoFisher | Cat# F10347 |
| Hoechst33342 | ThermoFisher | Cat# H3570 |
| Rapamycin | Sigma-Aldrich | Cat# R8781 |
| AZD-8055 | Nordic Bioste | Cat# HY-10422 |
| cOmplete™ EDTA-free Protease Inhibitor<br>Cocktail | Roche | Cat# 4693159001 |
| PhosphoStop inhibitor EDTA-free | Roche/Merck | Cat# 4906837001 |
| PageRuler™ Plus Prestained Protein<br>Ladder, 10 to 250 kDa | ThermoFisher | Cat# 26619 |
| Critical commercial assays |  |  |
| Click-iT EdU Cell Proliferation Kit | ThermoFisher | Cat# C10337/C10340 |
| Pierce™ Bradford Protein Assay Kit | ThermoFisher | Cat# 23246 |
| Clarity™ Western ECL Substrate | Bio-Rad | Cat# 1705060 |
| Deposited data |  |  |
| Flow cytometry files (.fcs) | This study | Deposited to FigShare |
| Western blot membrane raw images | This study | Deposited to Data Mendeley, and Table S8 |
| Experimental models: Organisms/strains |  |  |
| *Nematostella vectensis*: mOr2-Piwi1 transgenic line | Michael Sars Center | N/A |
| *Nematostella vectensis*: Piwi1-P2A-GFP transgenic line | Michael Sars Center | N/A |
| Software and algorithms |  |  |
| ImageJ/Fiji | Schindelin et al. | https://imagej.net/software/fiji/ |
| FlowJoV10.9 | BD Life Sciences | N/A |
| R v4.2.2 | N/A | https://www.r-project.org/ |

### Experimental model and subject details

#### Nematostella culture

*Nematostella vectensis* wildtype and transgenic polyps all derived from an original culture of CH6 females and CH2 males (Hand and Uhlinger, 1992). The genotype of transgenic polyps consisted of heterozygotes that resulted from a cross between homozygous *piwi1^mOr2^* or *piwi1^P2A-GFP^* knock-in lines (Miramón-Puértolas et al., 2024) and wildtype animals from the original stock.

Adult females and males were maintained in *Nematostella* medium (NM) at a salinity of 16‰, a temperature of 18°C and under dark conditions. Adults were fed fresh *Artemia nauplii* 5 times per week, with a daily, partial exchange of water and a full cleaning of the culture boxes once a month. Spawning was induced approximately every three weeks by 12-hour light exposure combined with a temperature shift from 18°C to 25 °C (Fritzenwanker and Technau, 2002; Hand and Uhlinger, 1992). Embryos were raised at 25°C and fed with mashed *Artemia nauplii* during the first week after reaching the polyp stage (∼10 days post-fertilization), followed by feeding with live *Artemia*. Sex could not be determined in juveniles therefore all data shown included both males and females. All experiments were performed on juvenile polyps of ∼5 mm in length just prior to the appearance of the 2nd pair of mesenteries.

### Method details

#### Feeding procedures

Polyps were starved for 5 or 20 days, representing short or long starvation periods. At these time points, they were either kept starved or refed *ad libitum* or for 1 hour. During starvation periods, dishes were cleaned using cotton swabs and NM was replaced two times per week. After 1-hour feedings, polyps were transferred to a new petri dish and NM was changed twice per week. Under *ad libitum* feeding conditions, polyps were fed twice and transferred to clean dishes daily.

To facilitate sampling, the 1-hour refeeding time point was delayed by 3, 6 and 9 hours, respectively (21/45-, 18/42- and 15/39-hours post-feeding samples). To assess potential circadian rhythmicity in cell proliferation dynamics, the T_0_+9 hours time point was sampled for each starvation condition.

#### EdU labelling

*Nematostella* polyps were relaxed with 0.1 M MgCl_2_ in NM before being transferred to NM containing 100 µM EdU (Invitrogen), 2% DMSO and 0.1 M MgCl_2_, or 2% DMSO (controls). Animals were incubated at room temperature (RT) for 30 minutes for a short EdU pulse and washed with 0.1 M MgCl_2_ in NM. In continuous EdU pulse experiments, animals were incubated at 25°C without MgCl_2_, and the EdU or DMSO solution was replaced every 24 hours. Flow cytometry analysis was performed after Trypsin/formaldehyde-based cell dissociation and fixation (see below). For microscopy sample preparation, polyps were fixed overnight in 3.7% formaldehyde/1× PBS, washed three times with 1× PBS/0.5% Triton X-100 and stored in 100% methanol (see below).

#### Trypsin/formaldehyde-based cell dissociation and fixation

Juveniles were dissociated in pools of 15 polyps following the published protocol (Garschall et al., 2024), with the resulting cell suspensions treated as biological replicates. Briefly, polyps were relaxed in 0.1 M MgCl_2_, washed with Ca^2+^- and Mg^2+^- free *Nematostella* medium (CMF/NM) followed by CMF/NM containing 0.195% ethylenediaminetetraacetic acid (CMF/NM+E). Then, the polyps were incubated for 5 minutes at 37°C in preheated CMF/NM+E containing 0.25% Trypsin (w/v). Animals were dissociated by pipetting and cold CMF/NM containing 1% BSA and 2.5% of Fetal Bovine Serum was added to stop trypsinization. Cells were pelleted at 800 g for 5 minutes at 4°C and resuspended in 1% BSA/1x PBS (w/v). After filtering through a pre-wetted 50 µm CellTrics strainer (Sysmex) the cell suspension was fixed with 3.7% formaldehyde for 30 minutes at RT in the dark. Finally, the cells were washed twice with 1% BSA/PBS by spinning at 800 g for 5 minutes at 4°C. The final cell pellet was resuspended in 90% Methanol/0.1%BSA/0.1xPBS in H_2_O and stored at -20°C. Before use, cell suspensions were rehydrated by washing twice with cold 1%BSA/PBS (800 g for 5 minutes at 4°C) and stained for flow cytometry.

#### S-phase detection and immunofluorescence on cell suspensions

After rehydration (see above), cells were permeabilized with 0.2% Triton X-100 in PBS for 15 minutes at RT and washed with 1X PBS. The cell pellet was resuspended in 50 µl freshly prepared Click-iT™ reaction cocktail containing Alexa488 (ThermoFisher, C10337) or Alexa647 fluorophore azide (ThermoFisher, C10340) for 30 minutes at RT in the dark, following manufacturer’s instructions. Cells were washed twice with 0.2% Triton X-100 in PBS (800 g for 5 minutes at 4°C). Cell suspensions were stained by immunofluorescence as previously described (Miramón-Puértolas et al., 2024). In short, cells were blocked in 1x PBS/10% DMSO/5% NGS/0.2% Triton X-100 for 30 minutes at RT. Primary antibody in 1x PBS/0.1% DMSO/5% NGS/0.2% Triton X-100 overnight at 4°C in the dark. The primary antibodies used were rabbit anti-DsRed 1:500 (Takara Bio Clontech 632496), mouse anti-mCherry 1:100 (Takara/AH Diagnostics 632543-CLO), rabbit anti-p-RPS6 (Ser235/236) 1:500 (Cell Signaling Technology, 4858) and rabbit anti-pH3 (Ser10) 1:500 (Cell Signaling Technology, 9701). After two washes with 1x PBS/0.2% Triton X-100 (800 g for 5 minutes at 4°C), cell suspensions were incubated with the secondary antibody in 1x PBS/0.1% DMSO/5% NGS/0.2% Triton X-100 for 30 minutes at RT in the dark. The secondary antibodies used were goat-anti-rabbit-Alexa488/568 1:500 (LifeTech, A-11008, A11011) and goat-anti-mouse-Alexa568 1:500 (LifeTech, A11011). Negative controls were stained with the secondary antibody only. Finally, cells were washed twice with 1x PBS/0.2% Triton X-100 (800 g for 5 minutes at 4°C) and resuspended in 1% BSA/PBS containing 1 µg/ml FXCycle Violet (ThermoFisher, F10347) for DNA staining. Cells were stored at 4°C and analyzed by flow cytometry within 24 hours without further washing.

#### Flow cytometry analysis

Flow cytometry was performed on a LSRFortessa^TM^ cell analyzer (BD Life Sciences) equipped with 407 nm, 488 nm, 561nm and 640 nm lasers. Cell cycle phases were distinguished using FXCycle violet DNA dye, detected with the BF450/50 filter. Incorporated EdU was coupled with Alexa647 fluorophore azide and detected using the BF670/14 filter. mOr-Piwi1 protein was detected with antibodies against either dsRed or mCherry coupled with Alexa568 and analyzed with the BF610/20 filter. phosphorylated Histone 3 and phosphorylated RPS6 were detected using specific pH3 and pRPS6 antibodies, respectively, coupled with Alexa488 and measured with the BF530/30 filter. The resulting data were analyzed using FlowJoV10.9 (BD Life Sciences). Graphical representations of all gating strategies are shown in Figure S1, S2, S6 and S7.

For gating cells of interest, debris was excluded based on size and granularity using the FSC-A/SSC-A gate, followed by FSC-A/FSC-H to exclude doublets and FSC-A/SSC-W to remove high-complexity events. DNA dye intensity was then gated by area over height to refine event selection. A histogram of DNA dye intensity (area, linear scale) was used to identify characteristic peaks corresponding to 2N and 4N DNA content. These pre-selected events constituted the pool of cells used for downstream analyses such as cell cycle composition (subfractions based on DNA signal intensity) and the fractions of EdU+, pRPS6+, pH3+ and mOr2-Piwi1+ cells, which were determined by gating fluorescence relative to negative controls. As previously described, we distinguished ‘low’ and ‘high’ mOr2 signal cells in the mOr2-Piwi1 population based on intensity and cell abundance. Throughout the manuscript, we referred to [mOr2-Piwi1]_high_ cells as mOr2-Piwi1+ or Vasa2+/Piwi1+ cells. The same gates were applied to analyze cell cycle composition and the fraction of EdU+, pRPS6+ and pH3+ cells within the population of mOr2-Piwi1+ cells. In the long-term EdU pulse, cell cycle composition was assessed based on DNA signal intensity without distinguishing S and G_2_/M populations.

#### S-phase detection and immunofluorescence on whole-mount tissues

Polyps were relaxed using 0.1 M MgCl_2_, fixed in 3.7% Formaldehyde NM for 1 hour at RT and the physa removed with a scalpel in a petri dish. The remaining polyp was washed thoroughly in 1x PBS/0.2% Tween20, followed by dehydration in a series of methanol washes (20-50-100% methanol in 1x PBS/0.2% Tween20) and in several 100% methanol washes until all pigment was washed out. Samples were stored in 100% methanol at -20°C. When needed, tissues were progressively rehydrated in 1xPBS/0.2% Triton X-100. In S-phase labelling, incorporated EdU was ‘clicked’ to a fluorophore azide using the Click-iT™ EdU Cell Proliferation Kit for Imaging (Invitrogen, C10337), following the manufacturer’s protocol. After 30 minutes Click-it staining reaction, tissue pieces were washed with 1xPBS/0.2% Triton X-100. For immunofluorescence staining, tissue pieces were blocked in 1x PBS/10% DMSO/5% normal goat serum (NGS)/0.2% Triton X-100 for 2 hours at RT. Primary antibody incubation was performed in 0.1% DMSO/5% NGS/0.2% Triton X-100 at 4°C using the following antibodies: FluoTag®-X4 anti-GFP 1:250 (NanoTag Biotechnologies, N0304) overnight and rabbit anti-p-RPS6 Ser235/236 1:200 (Cell Signaling Technology, 4858) over 3 nights. After washing with 1x PBS/0.2% Triton X-100, tissue was blocked in 1x PBS/5% NGS/0.2% Triton X-100 for 30 minutes at RT. Nuclear staining with Hoechst33342 (ThermoFisher, H3570) and secondary antibody incubation were performed in 1x PBS/5% NGS/0.2% Triton X-100 overnight at 4°C. The secondary antibody used was goat-anti-rabbit-Alexa568 (LifeTech A11011, A21244). Finally, tissue pieces were washed thoroughly in 1xPBS/0.2% Triton X-100, mounted on slides (Electron Microscopy Sciences, 63418-11) in 80% glycerol and sealed under a coverslip (Menzel-Gläser 18x18mm) with clear nail polish.

#### Confocal imaging

Immunofluorescence whole-mount tissue pieces were imaged on an Olympus FLUOVIEW FV3000 confocal microscope (standard PMT detectors) with 40x oil-immersion lens objective). The maximum projections stacks were processed, cropped and adjusted for levels and color balance with ImageJ/Fiji (Schindelin et al., 2012; Schindelin et al., 2015).

#### Drug treatment

A previously published Rapamycin incubation protocol for *Nematostella* (Garschall et al., 2024; Ikmi et al., 2020) was adapted as follows: a 10 mM Rapamycin (Sigma-Aldrich, R8781) stock solution in 100% DMSO was diluted in NM to a final concentration of 4 μM or 20 μM. For AZD-8055, the protocol was adapted from a previously published protocol for *Exaiptasia* (Voss et al., 2023): a 10 mM AZD-8055 (Nordic Bioste, HY-10422) diluted in 100% DMSO was diluted in NM to a final concentration of 0.1 and 1 μM. For all drug treatments, polyps were placed in drug solution for 2 hours before feeding. Fresh *Artemia nauplii* were added for 1 hour while polyps remained exposed to the drug. Afterwards, polys were transferred into a fresh drug solution. Incubations occurred at 25°C in the dark, with solutions replaced daily. Rapamycin, AZD-8055 or 0.2% DMSO (control) treatments were conducted over 1 or 2 days. When combining Rapamycin treatment with a continuous EdU pulse, the same protocol was followed. After the feeding pulse, polyps were transferred into a solution containing both the drug and EdU.

#### Western blotting

Protein was extracted from a pool of 50 juvenile polyps. After relaxation using 0.1 M MgCl_2_, polyps were transferred to homogenization tubes (M-tubes, Miltenyi Biotec, 130-093-236) containing RIPA buffer (150 mM NaCl, 50 mM Tris pH 8.0, 1%NP40, 0.5%DOC, 0.1% SDS) supplemented with cOmplete™ EDTA-free Protease Inhibitor Cocktail (Roche, 4693159001) and PhosphoStop inhibitor EDTA-free (Roche/Merck, 4906837001). Samples were incubated on ice for 30 minutes with intermittent mechanical disruption using the ‘protein_01_01’ program on the gentleMACS™ tissue homogenizer (Miltenyi Biotec, 130-093-235). The resulting homogenate was pelleted at 14,000 g for 15 min at 4°C, and the supernatant was transferred to a new tube and stored at −80°C. Protein concentration was quantified using the Bradford Assay Kit (Thermo Fisher Scientific, 23246) following the manufacturer’s protocol. For SDS-PAGE, 20 µg of protein was mixed 4:1 with 4× Laemmli sample buffer (0.1 M Tris-HCl pH 6.8, 2%SDS, 20% Glycerol, 4% β-mercaptoethanol, 0.02% Bromophenol blue) and boiled for 5 min before loading. Proteins were resolved on 7.5% Mini-PROTEAN® TGX™ precast gels (Bio-Rad, 4568096) in running buffer (25 mM Tris-HCl, 192 mM Glycine, 0.1% SDS) at 100 V for ∼90 min. The 10-250 kDa PageRuler™ Plus pre-stained protein ladder (Thermo Fisher Scientific, 26619) was used as a standard. Proteins were transferred to PVDF membranes using Trans-Blot Turbo Mini 0.2 µm PVDF Transfer Pack (Bio-Rad, 1704156) on a Trans-Blot Turbo transfer system (Bio-Rad) with the ‘mixed molecular weight’ program. Membranes were washed with 1× PBS/0.1% Tween (PBT) several times and blocked with 5% milk powder in PBT (MPBT) at RT for 1 h. Blots were cut at the 35 kDa band as a reference and incubated overnight at 4°C with the following primary antibodies in MPBT: rabbit anti-p-RPS6 (Ser235/236) (1:5000; Cell Signaling Technology, 4858) and rabbit anti-Actin (1:2000; Sigma-Aldrich, A5060). Membranes were washed several times in PBT and incubated in goat anti-rabbit-HRP (1:10.000; Abcam, Ab9705) secondary antibody in MPBT at RT for 1 h. After further washes with TBT (20 mM TrisHCl, 150 mM NaCl, 0.1% Tween pH 7.6), signals were detected using Clarify ECL substrate (Bio-Rad, 1705060) and imaged with a ChemiDoc XRS+ (Bio-Rad). Quantifications were performed with ImageJ software by measuring pixel intensity per band and subtracting the background intensity. The pRPS6 protein signal was normalized to the Actin signal. Membranes are shown in Table S8.

### Quantification and statistical analysis

#### Data visualization

Data visualization and analysis were performed using Excel and R v4.2.2 statistical analysis software (https://www.r-project.org/, R Core Team, 2021) with the packages ggplot2 (v3.5.1, (Wickham, 2016)), dplyr (v1.1.4, (Wickham et al., 2023)), stats (v4.2.2, R Core Team 2022), pracma (v2.4.4 (Borchers and Borchers, 2019)), minpack.lm (v1.2.4, (Elzhov et al., 2016)), bbmle (v0.25.1, (Bolker and Bolker, 2017)) and MASS (v3.58.1, (Venables and Ripley, 2013)).

Bar plots show the proportion of cell cycle phases, with the standard deviation of the lowest phase shown at phase intersections. Box plots show proportion of mOr2-Piwi1+ cells, EdU index, pRPS6 index, pH3 index and western blot quantifications. The central line shows the median, while the bounds represent the 25th and 75th percentiles. Whiskers extend to the maxima within 1.5x the interquartile range above the upper quartile and to the minima within 1.5x the interquartile range below the lower quartile. Dots represent single replicate data points. Scatter plots show EdU index and pRPS6 index over time, each data point represents a biological replicate. The line represents the mean and the overlay represents the 95% confidence intervals. In model plots, dots represent biological replicates for each time point, lines show predicted values and shaded areas indicate 95% confidence intervals.

#### Statistics

Statistical analysis included one-way ANOVA to estimate the effects of assay parameters (factors) followed by Tukey’s honest significant difference (HSD) test to evaluate pairwise differences between experimental groups. For parametric data comparison, a two-tailed Student’s t-test (α=0.05) was used. Correlations between p-RPS6 and EdU+ signal were estimated using Pearson correlation combined with Kolmogorov-Smirnov test, assessing normal Gaussian distribution. Values for the cEdU index under starvation condition was used as an input into linear regression model. Values for the cEdU index under *ad libitum* and 1-hour feeding conditions were used as input into Gompertz, Logisitic and Richards growth models.

Gompertz model equation

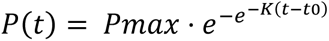

Logistic model equation

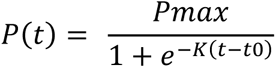

Richards model equation

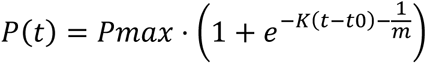

Akaike Information Criterion corrected (AICc) and coefficient of determination (R^2^) were calculated to support model selection. In sigmoidal growth models, steepness of the slope (K) and earlier half-max times (t50) obtained from Gompertz model indicate a faster response. Flow cytometry summary data and statistical analysis are provided in Table S1 - S7.

