## Supplementary Figures for "Nutritional regulation of cellular quiescence depth and cell cycle re-entry in Vasa2+/Piwi1+ cells in a sea anemone"

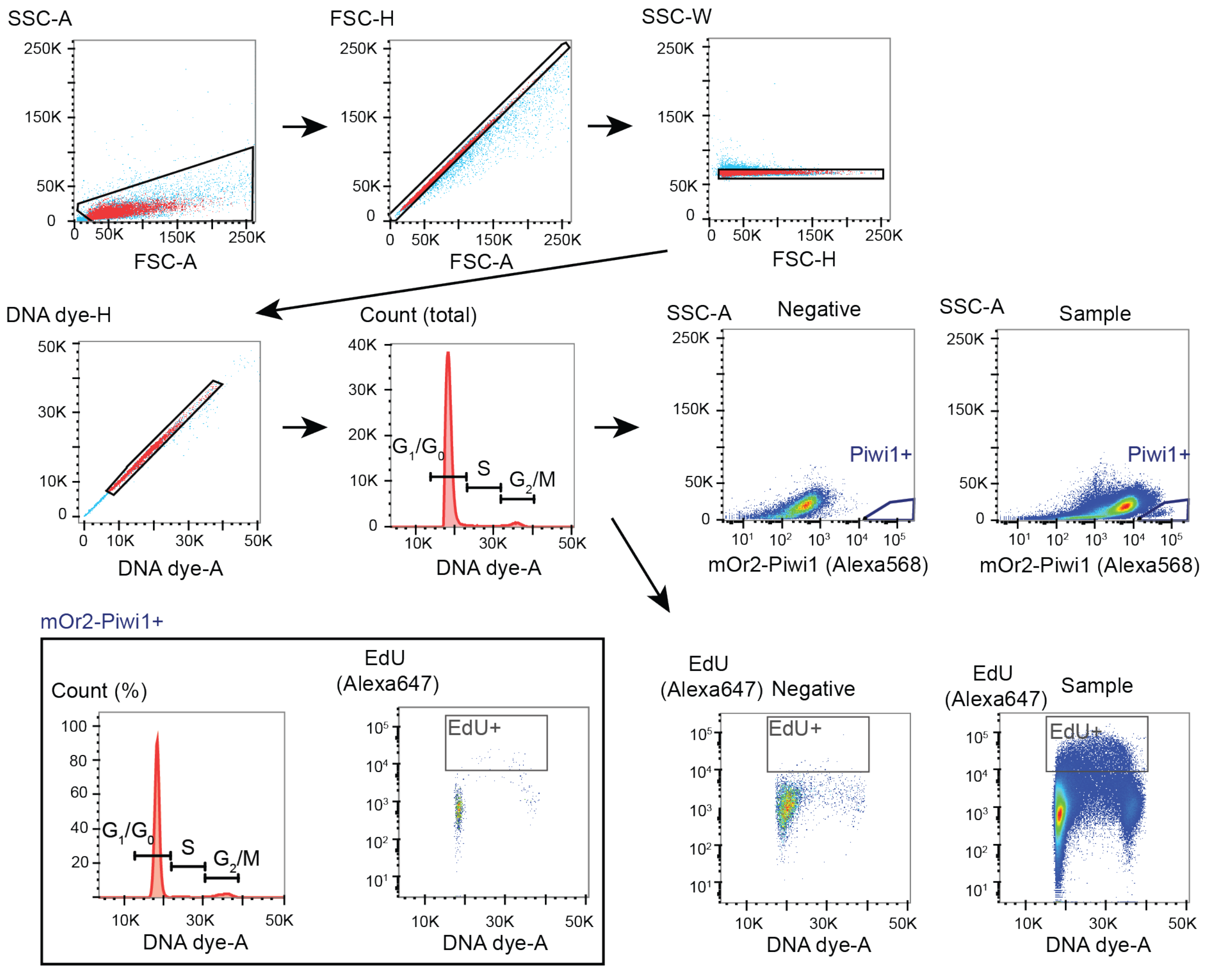


**Figure S1. Gating strategy of experiment using 30min EdU pulses experiments in mOr2-Piwi1 juvenile polyps. Related to Figure 2A-2F, 5E, 5F and S10E-S10G.**

Debris was excluded based on size and granularity in the FSC-A/SSC-A gate, with sub-gates based on FSC-A/FSC-H and FSC-A/SSC-W to remove potential cell doublets and high-complexity events. Then particles were gated based on DNA dye intensity in width-over-area plots, and a histogram of DNA dye intensity (area, linear scale) was created to visualize characteristic DNA peaks corresponding to cells between 2N and 4N. A threshold for EdU+ cells was determined based on the fluorescence signal of DMSO controls within the 2N–4N pool. Cells above this threshold were considered EdU+. Similarly, a threshold for mOr2-Piwi1+ cells was drawn based on the fluorescence signal of negative controls (no primary antibody) within the 2N–4N pool, identifying small and bright cells as mOr2-Piwi1+. Predefined cell cycle phases and EdU+ cells were then analyzed within this pool of cells.

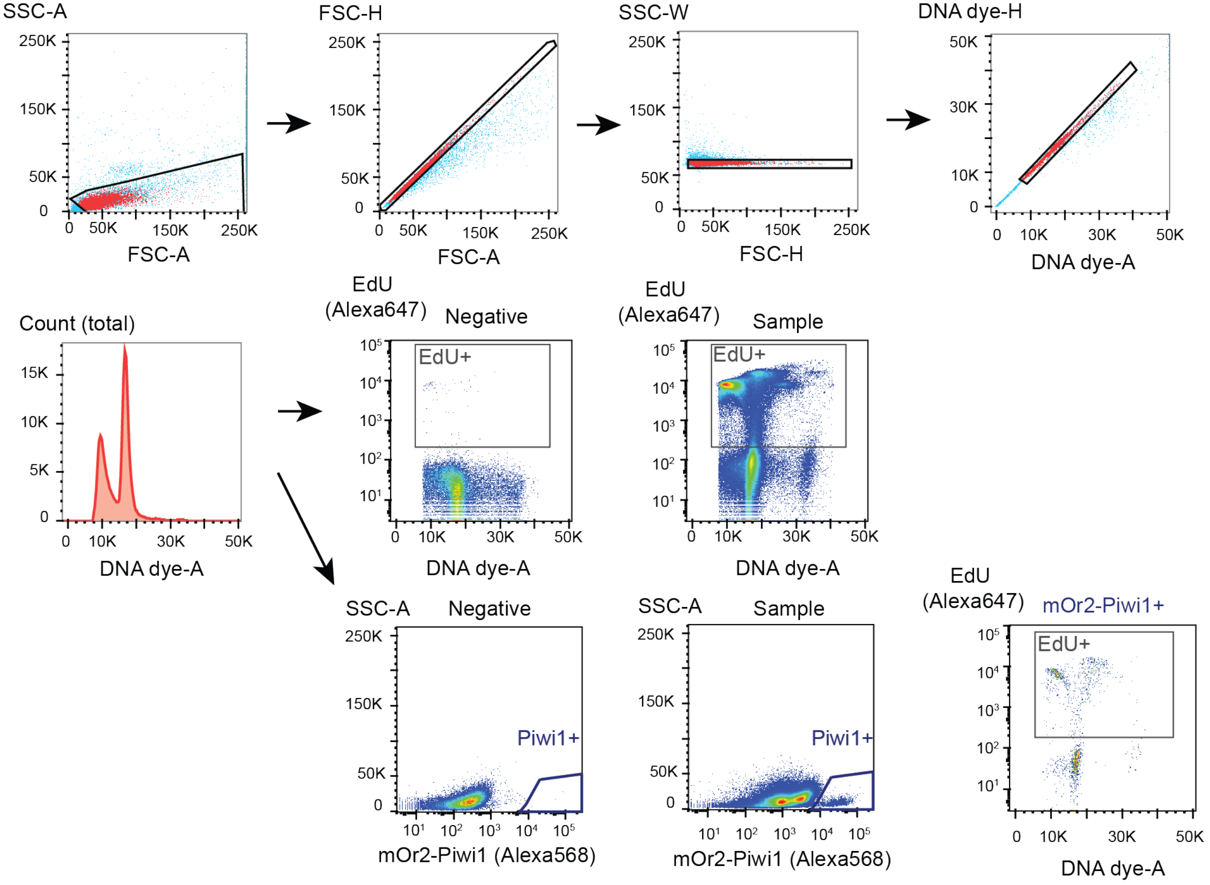


**Figure S2. Gating strategy of experiments using continuous EdU incubation in mOr2-Piwi1 juvenile polyps. Related to Figure 3A-3C, 5G, 5H, S3A-S3C, S4A and S4B.**

Debris was excluded based on size and granularity in the FSCA-/SSC-A gate, with sub-gates based in FSC-A/FSC-H and FSC-A/SSC-W to remove potential cell doublets and high-complexity events. Then, particles were gated based on DNA dye intensity in width-over-area plots, and a histogram of DNA dye intensity (area, linear scale) was created to visualize characteristic peaks corresponding to cells between 2N and 4N. We observed that long-term incorporation of EdU interfered with the DNA stain fluorescence and prevented a clear identification of 2N-4N cells. Therefore, we used a broader range of DNA intensity to define the parental gate of EdU+ populations. A threshold for EdU+ cells was determined based on the fluorescence signal of DMSO controls within the pool. Similarly, a threshold for mOr2-Piwi1+ cells was drawn based on the fluorescence signal of negative controls (no primary antibody), identifying small and bright cells as mOr2-Piwi1+. The same gates were applied to analyze cell cycle composition and the fraction of EdU+ cells within this pool of cells.

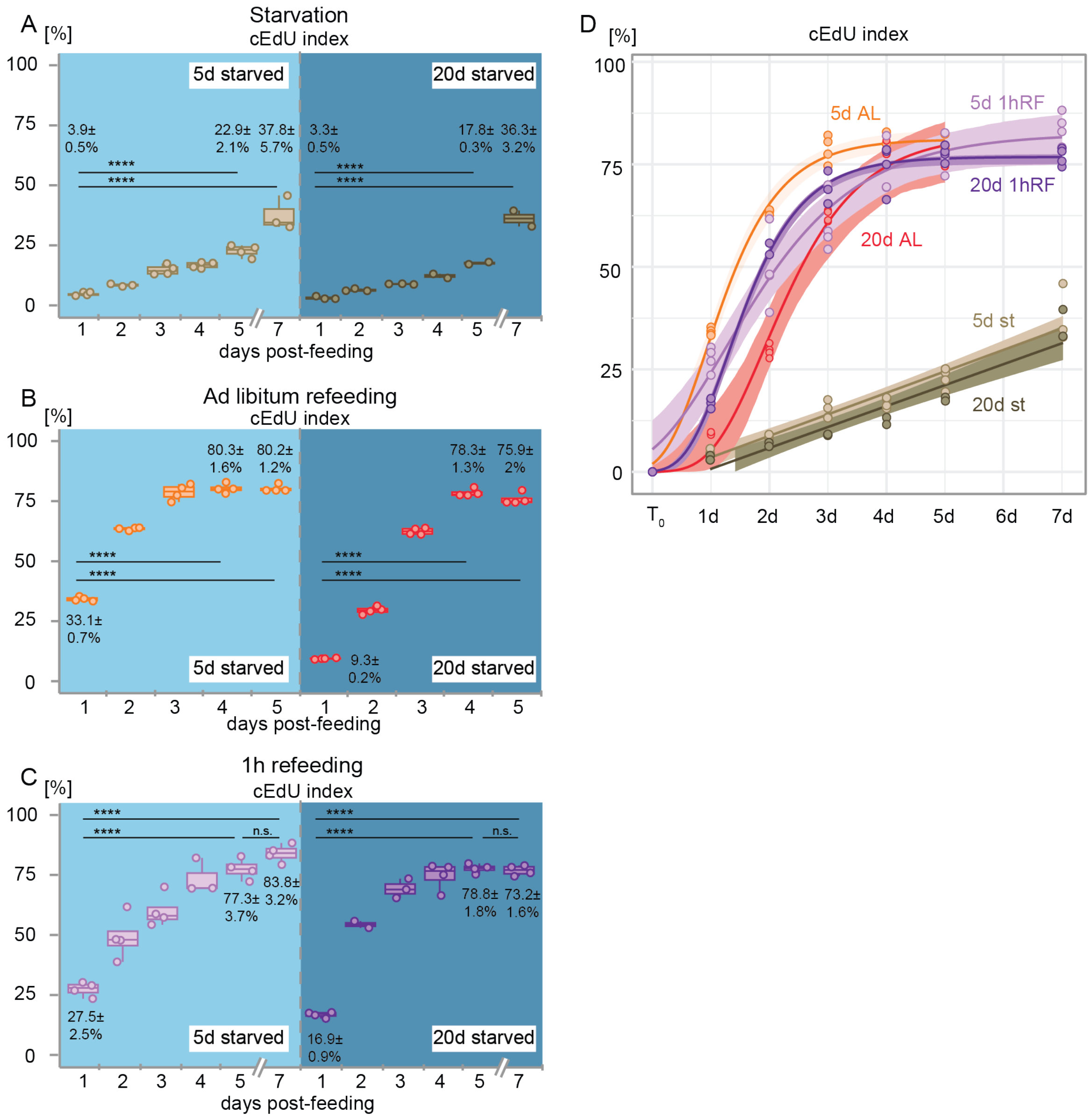


**Figure S3. The effect of feeding and starvation on the proliferative competence and onset of cell cycle re-entry in all cell cycle-gated cells. Related to Figure 3.**

(A-C) Temporal changes in the cumulative EdU (cEdU) index under continued starvation (A), *ad libitum* refeeding (B) or following a single, 1-hour refeeding pulse (C) after 5 or 20 days of starvation in all cells within the cell cycle. Experiments were done using flow cytometry. (D) Dynamics of the cEdU index (A-C) are best explained by linear growth models under continued starvation (st), or by Gompertz growth models after *ad libitum* (AL) or a 1-hour refeeding pulse (1hRF). Dots represent same replicate sample values as in (A-C). *n*=2-4 biological replicates per condition (15 individuals per replicate). Coloured lines in D represent the model curve or line for each condition with overlays depicting 95% confidence intervals. See 'Data visualisation' for definition of box plots. Dots represent individual values. Index values represent means ± standard deviations of respective timepoints. Pairwise comparisons after one-way ANOVA were calculated using Tukey's HSD and *p* values adjusted at significance codes: ****$p$<0.0001. d: day(s), n.s.: non-significant. See also Table S3 and S4.

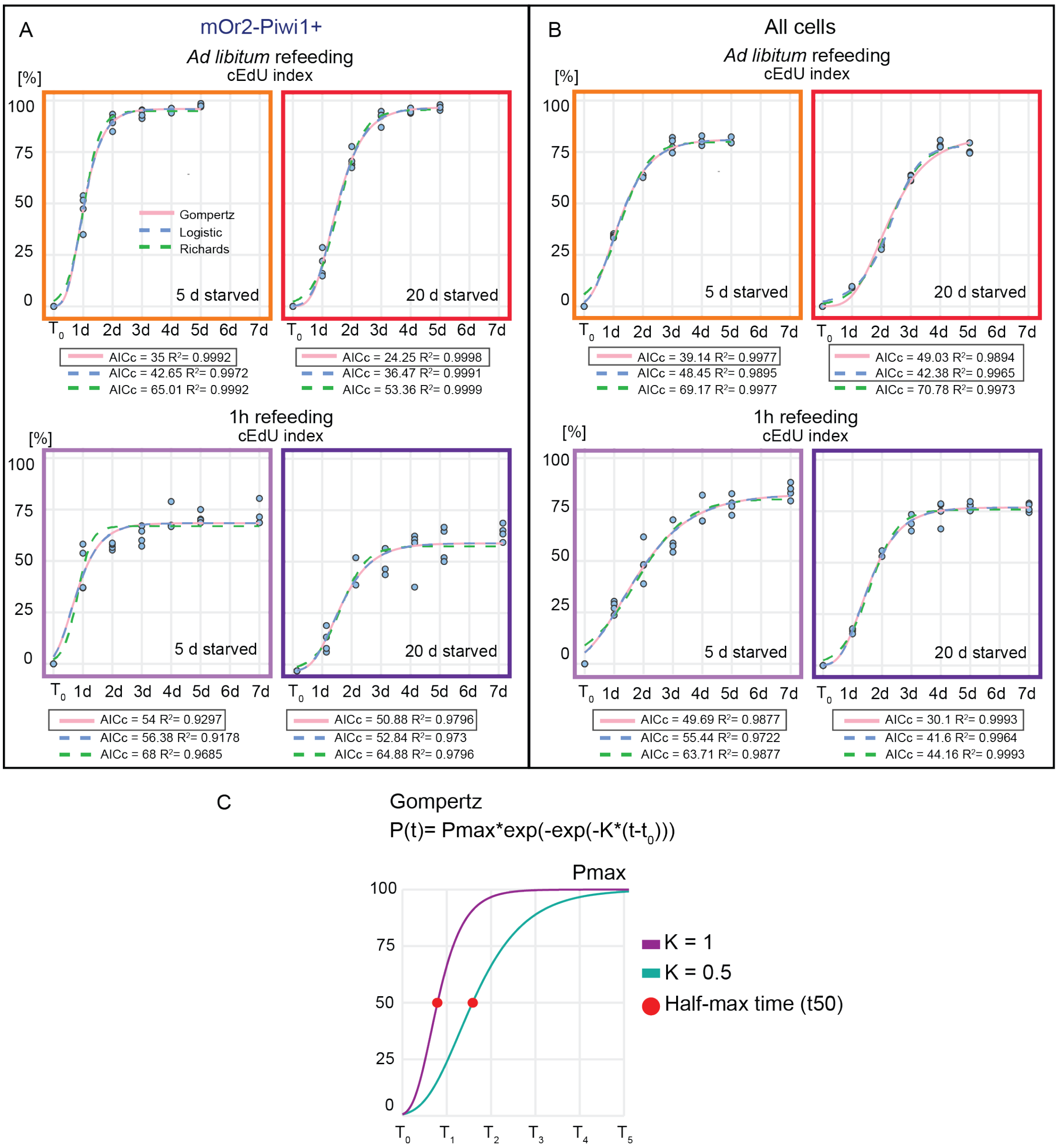


**Figure S4. Model selection to estimate the dynamics of EdU+ cell accumulation during *ad libitum* and 1-hour feeding conditions. Related to Figure 3.**

(A, B) Comparison of Gompertz (solid pink), Logistic (dashed blue) and Richards (dashed green) growth models for mOr2-Piwi1+ (A) and all cell cycle-gated cells (B) during *ad libitum* and 1-hour refeeding conditions. The best fitting model for each condition was chosen based on the highest coefficient of determination ($R^2$) and lowest Akaike Information Criterion corrected (AICc). Over all conditions, the Gompertz growth model performed best (dark boxes). *n*=2-4 biological replicates per condition (15 individuals per replicate). Coloured lines in (A, B) represent the model curve or line for each condition. Dots represent individual values. (C) The Gompertz growth model assumes a maximum value and exponential decay as the population approaches this maximum. The growth rate constant (K) controls how quickly the curve transitions and indicates the speed at which the predicted maximum (Pmax) is reached. The half-max time (t50) indicates the time point where the function reaches 50% of the Pmax. d: day(s). See also Table S4.

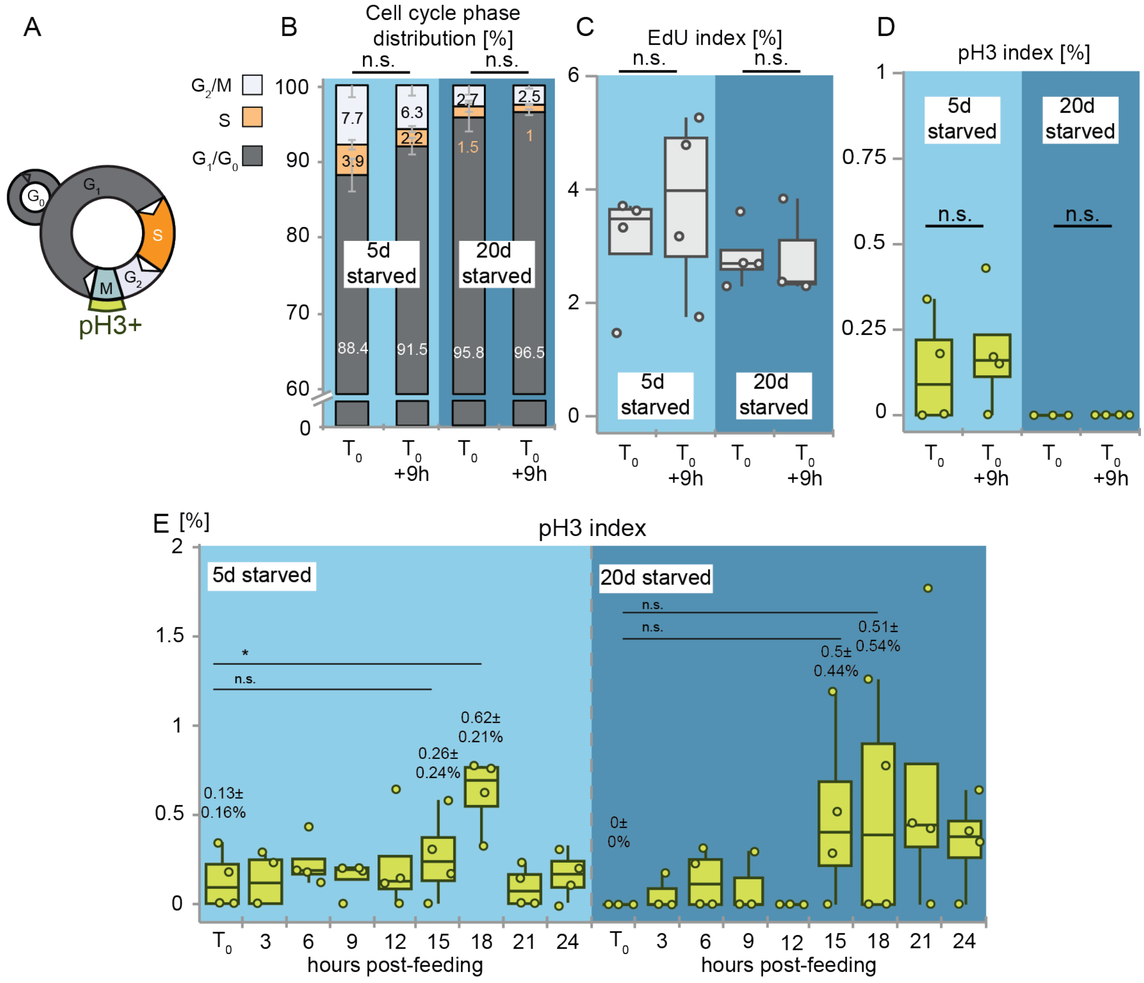


**Figure S5. Proliferation rates of Vasa2+/Piwi1+ were independent on the daytime of the sampling, and mitotic rates were not significantly different upon refeeding of 5 or 20 days starved polyps. Related to Figure 4.**

(A) Schematics of cell cycle phases, highlighting the phospho-histone H3 (pH3) labelling of the M-phase. (B-D) The cell cycle phase distribution (B), proportion of EdU+ cells (EdU index; C) and proportion of pH3+ cells (pH3 index; D) of Vasa2+/Piwi1+ cells sampled 9h apart showed no significant difference. (E) Quantification of the pH3 index over 24 hours after refeeding polyps starved for 5 or 20 days showed no significant difference between $T_0$ and 15h regardless of the starvation length. All experiments (B-D) were done using flow cytometry. For box plots and bar plot definitions, see 'Data visualisation'. Dots in (C-E) represent individual values. Values in (E) represent means ± standard deviations of respective timepoints with dots representing individual samples. *n*= 3-4 biological replicates per condition, with 15 polyps per replicate. Pairwise comparisons after one-way ANOVA were calculated using Tukey's HSD and *p* values adjusted at significance codes: $^*p<0.05$. n.s.: non-significant. See also Table S5.

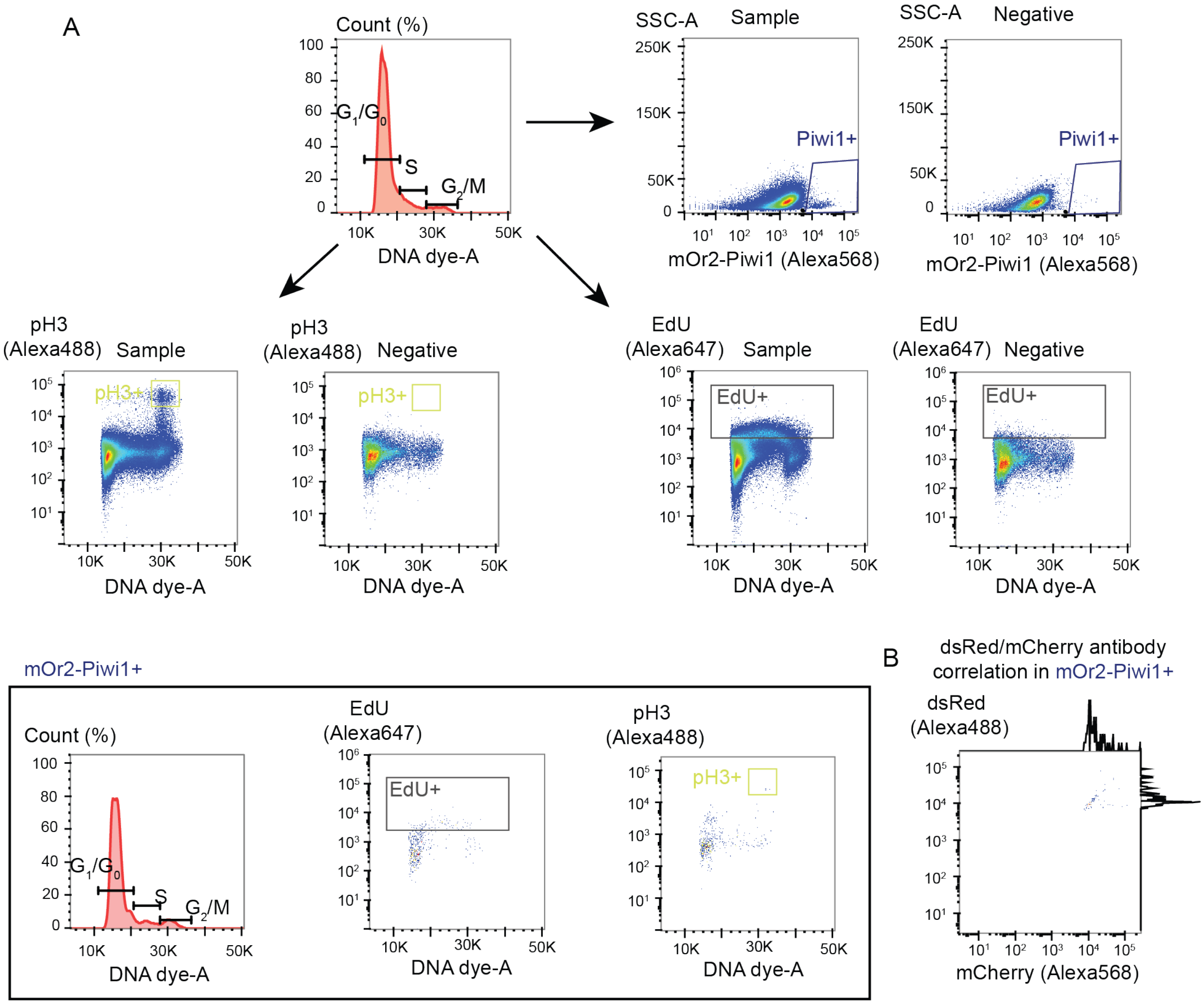


**Figure S6. Gating strategy of experiment using 30min EdU pulses and pH3 detection in mOr2-Piwi1 juvenile polyps. Related to experiments in Figure 4A, 4B and S5C-S5F.**

Debris was excluded, and cells were gated based on DNA dye intensity as described above. For EdU+ cells, a threshold was drawn above the fluorescence signal of DMSO controls within the 2N–4N pool. For pH3+ cells, a threshold was drawn based on the fluorescence signal of negative controls (no primary antibody) within the 2N–4N pool, identifying G2/M-phase cells as expected. Similarly, for mOr2-Piwi1+ cells, a threshold was drawn based on the fluorescence signal of negative controls within the 2N–4N pool, identifying a small population of bright cells. The same gates were applied to analyze cell cycle phases and the proportion of pH3+ and EdU+ cells within this pool of cells. (B) Comparison of mCherry and dsRed antibodies for the immunolabeling of mOr2-Piwi1 cells. mOr2-Piwi1 was detected using both an mCherry antibody coupled with Alexa568 and a dsRed antibody coupled with Alexa488. Debris was excluded, and cells were gated based on DNA-dye intensity as explained above. For mOr2-Piwi1+ cells, thresholds were drawn based on the fluorescence signal of negative controls (no primary antibody) within the 2N–4N pool. In this population of cells, a linear correlation between the fluorescent signals was confirmed.

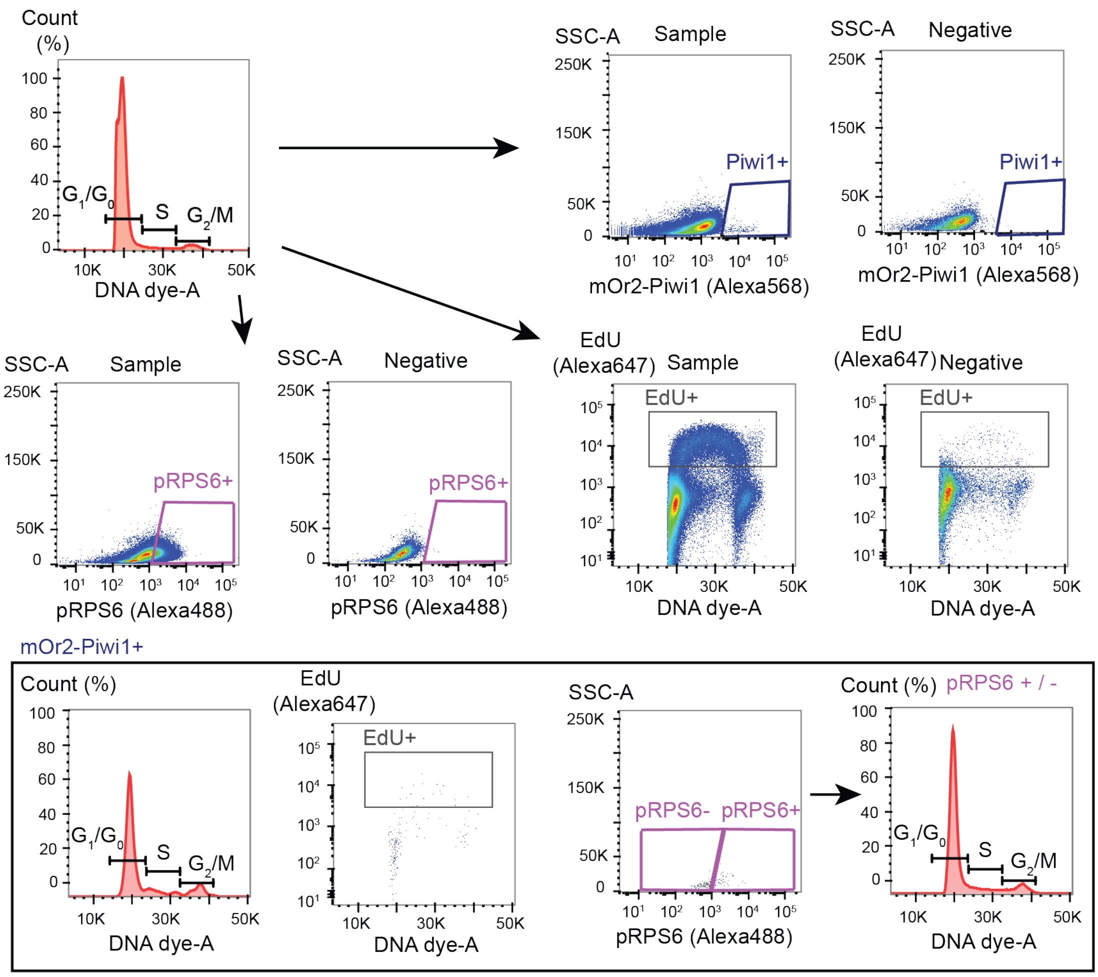


**Figure S7. Gating strategy in short EdU (30 min) and pRPS6 experiments in mOr2-Piwi1 juvenile polyps. Related to Figure 5A-5D, S8B-S8D and S9A-S9D.**

Debris was excluded and gated cells based on DNA-dye intensity as explained above. For EdU+ cells, a threshold was drawn above the fluorescence signal of DMSO controls within the 2N-4N pool of cells. For pRPS6+ and pRPS6- cells, thresholds were drawn based on the fluorescence signal of negative controls (no primary antibody) within 2N-4N pools of cells. For mOr2-Piwi1+ cells, a threshold was drawn based on the fluorescence signal of negative controls (no primary antibody) within 2N-4N pools of cells. The same gates were applied to analyze cell cycle phases and the proportion of pRPS6+ and EdU+ cells within this pool of cells.

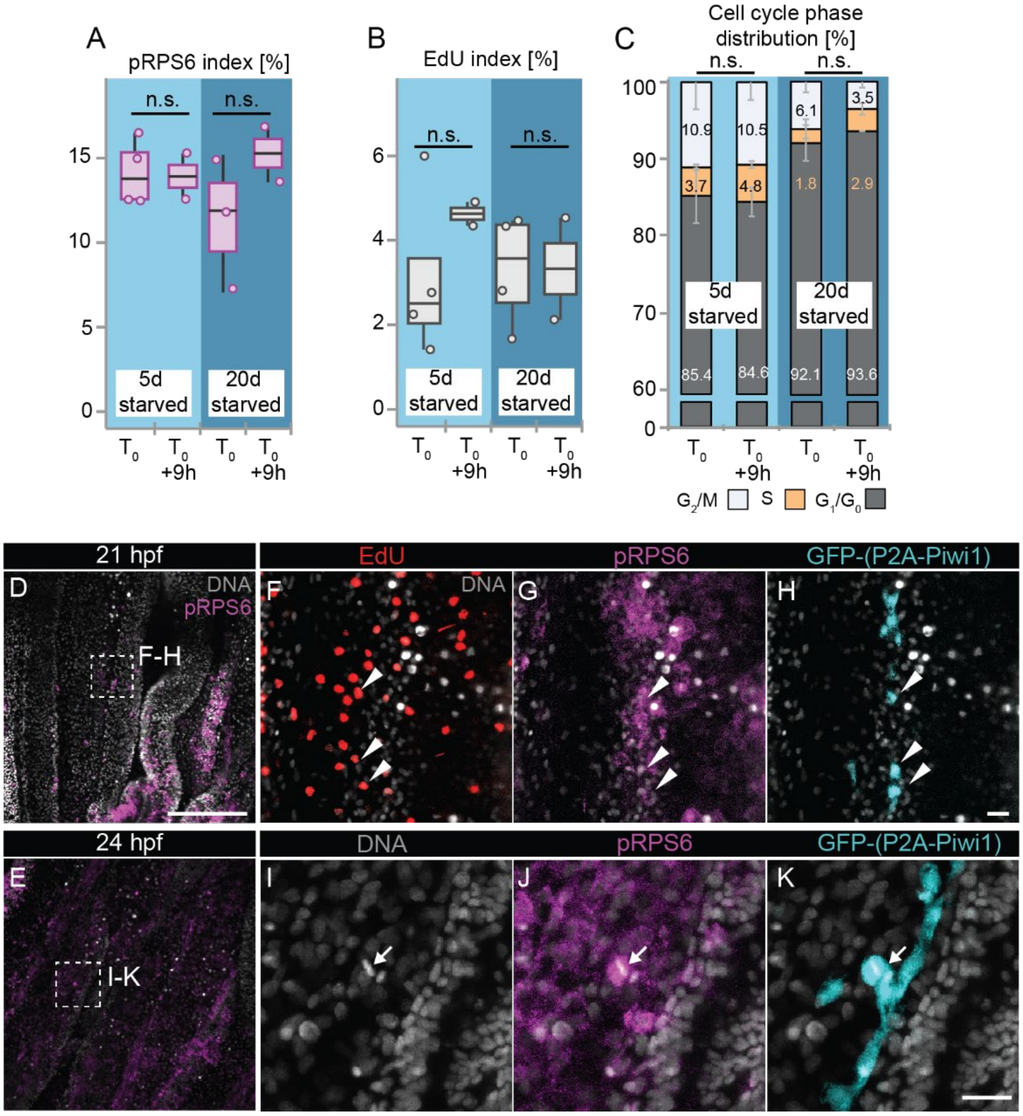


**Figure S8. pRPS6 and proliferation rates of Vasa2+/Piwi1+ cells were independent on the daytime of refeeding, and phospho-RPS6 was found in S- and M-phase Vasa2+/Piwi1+ cells. Related to Figure 5.**

(A-C) The proportion of phospho-ribosomal protein S6-positive cells (pRPS6 index; A), EdU+ cells (EdU index; B) and cell cycle phase distribution (C) of Vasa2+/Piwi1+ cells sampled 9h apart showed no significant differences. Experiments were done using flow cytometry. For box plots and bar plot definitions, see 'Data visualisation'. Dots in (A, B) represent individual values. *n*= 2-4 biological replicates per condition, with 15 polyps per replicate. (D-K) Confocal imaging stacks of gastrodermal tissue from Piwi1$^{P2A\text{-}GFP}$ juvenile polyps. (D, E) Overview of mesenteries at midbody level of whole-mount polyps stained by immunofluorescence against pRPS6 sampled at 21h or 24h post-refeeding (hpf) after 5 days of starvation. Side views with oral side oriented downwards. (F-H) Single cells co-labelled by EdU, pRPS6 and Piwi1-(P2A-GFP)(white arrowheads). (I-K) A single metaphase cell co-labelled by pRPS6 and Piwi1-(P2A-GFP)(white arrow). EdU pulse labelling started 30 minutes before fixation. Gray: Hoechst DNA dye. Scale bar: 100 μm (D, E) and 10 μm (F-K). n.s.: non-significant. See also Table S6.

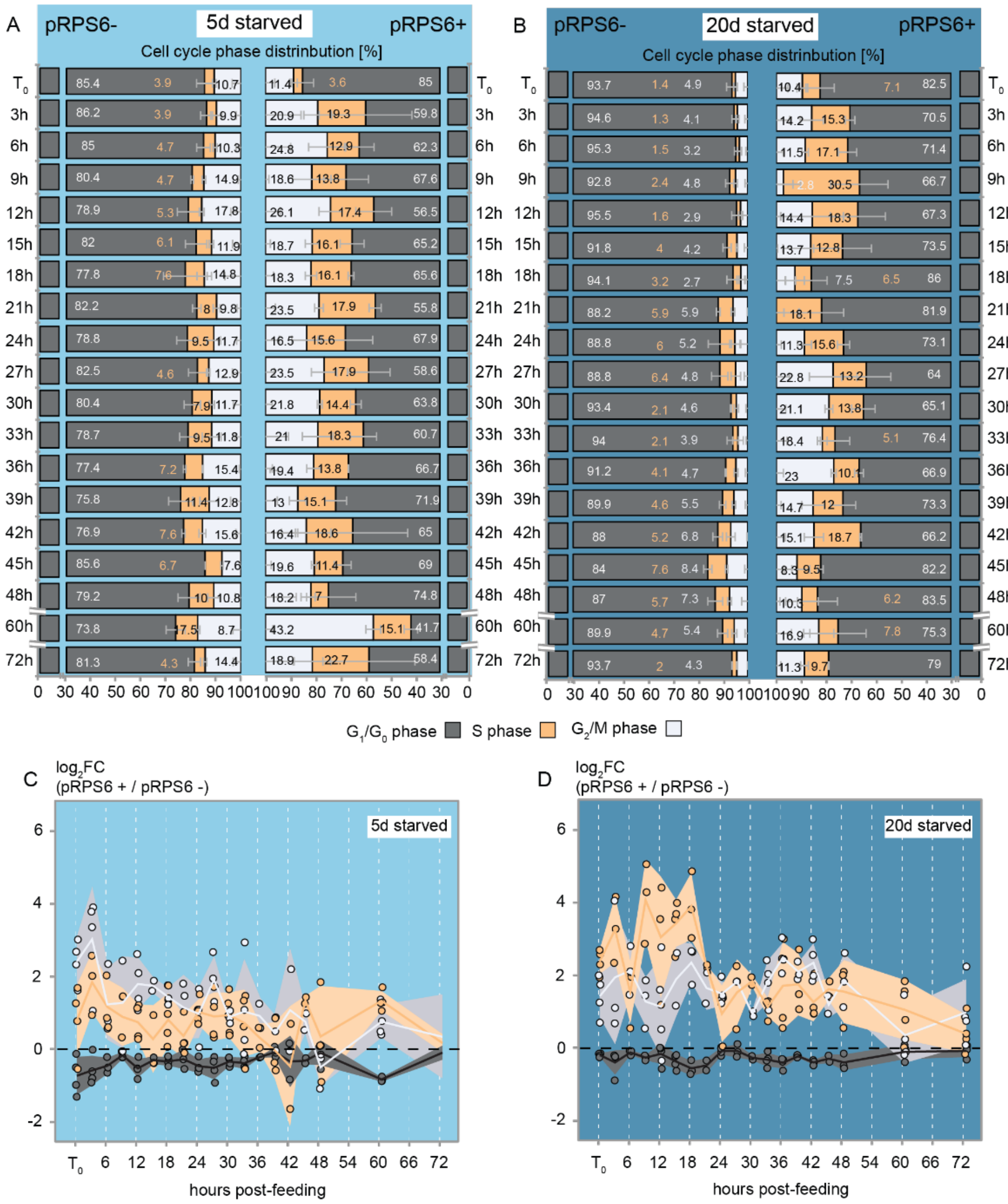


**Figure S9. Overrepresentation of S and G2/M cells within the phospho-RPS6+ fraction of Vasa2+/Piwi1+ cells. Related to Figure 5.**

(A, B) Flow cytometry-based cell cycle phase distributions of pRPS6+ and pRPS6– cells over 72 hours after refeeding of 5 (A) or 20 (B) days starved polyps. At $T_0$, polyps were refed for 1 hour and sampled at indicated time points. For bar plot definitions, see 'Data visualisation'. Values in (A, B) represent means. *n*= 2-4 biological replicates per condition, with 15 polyps per replicate. (C, D) $Log_2FC$ of the ratio of the cell cycle fractions between pRPS6+ and pRPS6– cells. Note that regardless of the starvation duration, the pRPS6+ cells are overrepresented ($Log_2FC>0$) in the S and $G_2/M$ fractions upon refeeding. Dots represent individual samples. Coloured lines indicate mean values for each cell cycle phase and band overlays represent 95% confidence intervals. *n*= 2-4 biological replicates per condition, with 15 polyps per replicate. See also Table S6.

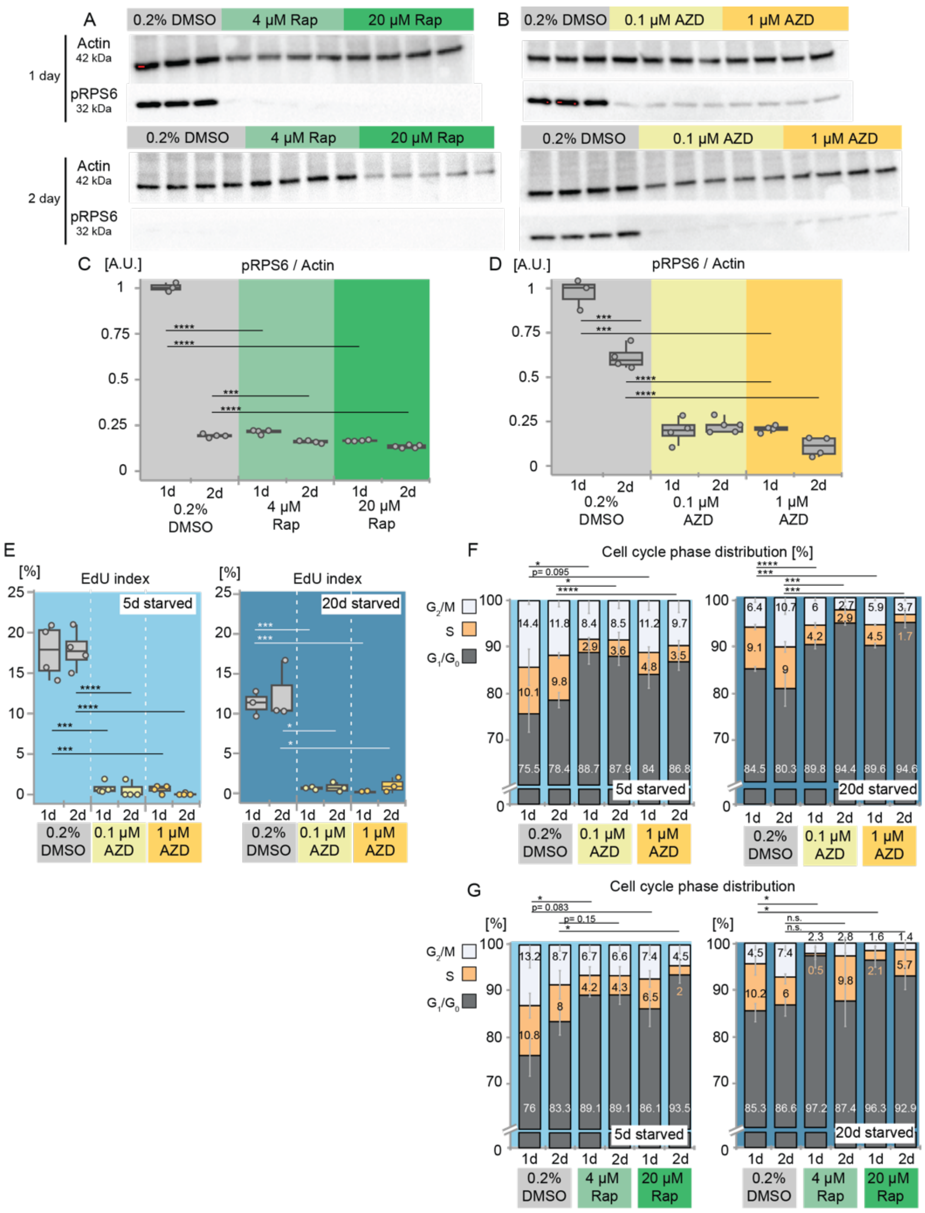


**Figure S10. The TOR inhibitors Rapamycin and AZD-8055 strongly reduce RPS6 phosphorylation and cell proliferation in Vasa2+/Piwi1+ cells. Related to Figure 5.**

(A, B) Western blots depict protein levels of phosphorylated ribosomal protein S6 (pRPS6) Actin (as control) after refeeding and incubating for 1 or 2 days with 0.2% DMSO, 4 µM or 20µM Rapamycin (‘Rap’, A) or 0.1 µM or 1µM AZD-8055 (‘AZD’, B). (C, D) Intensity measures of pRPS6 bands relative to the Actin control protein show that Rapamycin (C) and AZD-8055 (D) led to a decrease of phosphorylated RPS6 levels. *n*= 3-5 technical replicates from one biological replicate with pools of 50 Rapamycin-, AZD-8055- or 0,2% DMSO-treated polyps. (E-G) Effect of AZD-8055 (E, F) and Rapamycin (G) treatment on the proportion of EdU (EdU index, E) and cell cycle phase distribution (F, G) after a 1h-feeding pulse at 5 or 20 days of starvation. Compared to 0.2% DMSO-treated controls, AZD leads to a reduced EdU index regardless of starvation history, concentration or incubation time (E). AZD (F) and Rap (G) reduced the fractions of S- and $G_2$/M-phase cells. For box plots and bar plot definitions, see 'Data visualisation'. Dots represent individual values. *n*= 2-4 biological replicates per condition, with 15 individuals per replicate. Significance levels for Student's t-test are indicated for adjusted *p* values: *$p<0.05$, ***$p<0.001$, ***$p<0.0001$. d: day(s), n.s.: non-significant. See also Table S7.

### Table S1. Related to Figure 2.

**Table S1A.** Flow cytometer analysis after 30min of EdU pulse - cell cycle phases defined as per DNA content

**Table S1B.** Flow cytometer analysis after 30min of EdU pulse - cell cycle phases defined as per DNA content - extended starvation

**Table S1C.** ANOVA for the effect of **feeding and starvation day** on the fraction of **mOr2-Piwi1 high** and pairwise comparisons between days (Tukey's HSD)

**Table S1D.** ANOVA for the effect of **feeding and starvation day** on the **EdU index** and pairwise comparisons between days (Tukey's HSD)

**Table S1E.** ANOVA for the effect of **feeding and starvation day** on the fraction of **S-phase cells** and pairwise comparisons between days (Tukey's HSD)

**Table S1F.** ANOVA for the effect of **starvation day** on the fraction of **mOr2-Piwi1 high** and pairwise comparisons between days (Tukey's HSD)

**Table S1G.** ANOVA for the effect of **starvation day** on the **EdU index** and pairwise comparisons between days (Tukey's HSD)

**Table S1H.** ANOVA for the effect of **starvation day** on the fraction of **S-phase cells** and pairwise comparisons between days (Tukey's HSD)

### Table S2. Related to Figure 3 and S3.

**Table S2A.** Flow cytometer analysis after continuous EdU pulse during starvation - cell cycle phases defined as per DNA content in mOr2-Piwi1 cells

**Table S2B. 5 days starved** - ANOVA for the effect of **starvation day** on the **cEdU index** and pairwise comparisons between days (Tukey's HSD) - **mOr2-Piwi1 cells**

**Table S2C. 20 days starved** - ANOVA for the effect of **starvation day** on the **cEdU index** and pairwise comparisons between days (Tukey's HSD) - **mOr2-Piwi1 cells**

**Table S2D.** Flow cytometer analysis after continuous EdU pulse during *ad libitum* - cell cycle phases defined as per DNA content in mOr2-Piwi1 cells

**Table S2E. 5 days starved** - ANOVA for the effect of ***ad libitum*** **day** on the **cEdU index** and pairwise comparisons between days (Tukey's HSD) - **mOr2-Piwi1 cells**

**Table S2F. 20 days starved** - ANOVA for the effect of ***ad libitum*** **day** on the **cEdU index** and pairwise comparisons between days (Tukey's HSD) - **mOr2-Piwi1 cells**

**Table S2G.** Flow cytometer analysis after continuous EdU pulse after 1h Refed - cell cycle phases defined as per DNA content in mOr2-Piwi1 cells

**Table S2H. 5 days starved** - ANOVA for the effect of **1h Refed day** on the **cEdU index** and pairwise comparisons between days (Tukey's HSD) - **mOr2-Piwi1 cells**

**Table S2I. 20 days starved** - ANOVA for the effect of **1h Refed day** on the **cEdU index** and pairwise comparisons between days (Tukey's HSD) - **mOr2-Piwi1 cells**

**Table S2J. DAY 1** - ANOVA for the effect of **feeding regime** on the **cEdU index** and pairwise comparisons between regimes (Tukey's HSD) in **mOr2-Piwi1 cells**

**Table S2K. DAY 5 -** ANOVA for the effect of **feeding regime** on the **cEdU index** and pairwise comparisons between regimes (Tukey's HSD) in **mOr2-Piwi1 cells**

**Table S2L. DAY 7 -** ANOVA for the effect of **feeding regime** on the **cEdU index** and pairwise comparisons between regimes (Tukey's HSD) in **mOr2-Piwi1 cells**

### Table S3. Related to Figure S3.

**Table S3A.** Flow cytometer analysis after continuous of EdU pulse during starvation - cell cycle phases defined as per DNA content in All cells

**Table S3B. 5 days starved** - ANOVA for the effect of **starvation day** on the **cEdU index** and pairwise comparisons between days (Tukey's HSD) - **All cells**

**Table S3C. 20 days starved** - ANOVA for the effect of **starvation day** on the **cEdU index** and pairwise comparisons between days (Tukey's HSD) - **All cells**

**Table S3D.** Flow cytometer analysis after continuous of EdU pulse during *ad libitum* - cell cycle phases defined as per DNA content in All cells

**Table S3E. 5 days starved** - ANOVA for the effect of ***ad libitum*** **day** on the **cEdU index** and pairwise comparisons between days (Tukey's HSD) - **All cells**

**Table S3F. 20 days starved** - ANOVA for the effect of ***ad libitum*** **day** on the **cEdU index** and pairwise comparisons between days (Tukey's HSD) - **All cells**

**Table S3G.** Flow cytometer analysis after continuous of EdU pulse after 1h Refed - cell cycle phases defined as per DNA content in All cells

**Table S3H. 5 days starved** - ANOVA for the effect of **1h Refed day** on the **cEdU index** and pairwise comparisons between days (Tukey's HSD) - **All cells**

**Table S3I. 20 days starved** - ANOVA for the effect of **1h Refed day** on the **cEdU index** and pairwise comparisons between days (Tukey's HSD) - **All cells**

**Table S3J. DAY 1** - ANOVA for the effect of **feeding and starvation day** on the **cEdU index** and pairwise comparisons between days (Tukey's HSD) in **mOr2-Piwi1 cells**

**Table S3K. DAY 5 - ANOVA for the effect of feeding and starvation day** on the **cEdU index** and pairwise comparisons between days (Tukey's HSD) in **mOr2-Piwi1 cells**

**Table S3L. DAY 7 - ANOVA for the effect of feeding and starvation day** on the **cEdU index** and pairwise comparisons between days (Tukey's HSD) in **mOr2-Piwi1 cells**

### Table S4. Related to Figure 3, S3 and S4.

**Table S4A. Linear regression** analysis of EdU+ cell accumulation

**Table S4B. Growth model analysis** of EdU+ cell accumulation

**Table S4C. Gompertz growth** model analysis of EdU+ cell accumulation: **K** and **t50**

### Table S5. Related to Figure 4 and S5.

**Table S5A.** Flow cytometer analysis after 30min of EdU pulse - cell cycle phases defined as per DNA content - pH3 index

**Table S5B. 5/20 days starved** - ANOVA for the effect of **starvation day** on the fraction of **S-phase cells, G2/M-phase cells EdU index and pH3 index** and pairwise comparisons between **T0 and T0+9h** (Tukey's HSD)

**Table S5C. 5 days starved** - ANOVA for the effect of **feeding and starvation day** on the **EdU index** and pairwise comparisons between days (Tukey's HSD)

**Table S5D. 5 days starved** - ANOVA for the effect of **feeding and starvation day** on the fraction of **S-phase cells** and pairwise comparisons between days (Tukey's HSD)

**Table S5E. 5 days starved** - ANOVA for the effect of **feeding and starvation day** on the **pH3 index** and pairwise comparisons between days (Tukey's HSD)

**Table S5F. 20 days starved** - ANOVA for the effect of **feeding and starvation day** on the **EdU index** and pairwise comparisons between days (Tukey's HSD)

**Table S5G. 20 days starved** - ANOVA for the effect of **feeding and starvation day** on the fraction of **S-phase cells** and pairwise comparisons between days (Tukey's HSD)

**Table S5H. 20 days starved** - ANOVA for the effect of **feeding and starvation day** on the **pH3 index** and pairwise comparisons between days (Tukey's HSD)

### Table S6. Related to Figure 5, S8 and S9.

**Table S6A.** Flow cytometer analysis after 30min of EdU pulse - cell cycle phases defined as per DNA content - pRPS6 index

**Table S6B. 5/20 days starved** - ANOVA for the effect of **starvation day** on the fraction of **S-phase cells, EdU index and pRPS6 index** and pairwise comparisons between **T0 and T0+9h** (Tukey's HSD)

**Table S6C.** Flow cytometer analysis after 30min of EdU pulse - pRPS6 index

**Table S6D. 5 days starved** - **Pearson correlation** between EdU+ and pRPS6+ cells

**Table S6E. 20 days starved** - **Pearson correlation** between EdU+ and pRPS6+ cells

**Table S6F. 20 days starved_$T_0$ to 30hpf** - **Pearson correlation** between EdU+ and pRPS6+ cells

**Table S6G. 20 days starved_30hpf to 72hpf** - **Pearson correlation** between EdU+ and pRPS6+ cells

**Table S6H.** Flow cytometer analysis cell cycle phases defined as per DNA content, of pRPS6+ and pRPS6- cells

**Table S6I.** Log2FC pRPS6+ / pRPS6- Cell cycle phases

### Table S7. Related to Figure 5 and S10.

**Table S7A.** Western blot analysis after Rapamycin and AZD-8055 treatments - pRPS6/Actin levels

**Table S7B.** Student's t-test pairwise comparisons between relative **pRPS6/Actin levels** from **Rapamycin** treated samples and controls

**Table S7C.** Student's t-test pairwise comparisons between relative **pRPS6/Actin levels** from **AZD-8055** treated samples and controls

**Table S7D.** Rapamycin treatment - Flow cytometer analysis after 30min of EdU pulse - cell cycle phases defined as per DNA content

**Table S7E. 5/20 days starved** - Student's t-test pairwise comparisons between the **EdU index** from **Rapamycin** treated samples and controls with **short EdU pulse**

**Table S7F. 5/20 days starved** - Student's t-test pairwise comparisons between the fraction of **S-phase cells** from **Rapamycin** treated samples and controls with **short EdU pulse**

**Table S7G.** AZD-8055 treatment - Flow cytometer analysis after 30min of EdU pulse - cell cycle phases defined as per DNA content

**Table S7H. 5/20 days starved** - Student's t-test pairwise comparisons between the **EdU index** from **AZD-8055** treated samples and controls with **short EdU pulse**

**Table S7I. 5/20 days starved** - Student's t-test pairwise comparisons between the fraction of **S-phase cells** from **AZD-8055** treated samples and controls with **short EdU pulse**

**Table S7J.** Rapamycin treatment - Flow cytometer analysis after continuous EdU pulse - cell cycle phases defined as per DNA content

**Table S7K. 5/20 days starved** - Student's t-test pairwise comparisons between the **cEdU index** from **Rapamycin** treated samples and controls with **continuous EdU pulse**

### Table S8. Related to Figure S10. Western blot membranes
